# Influence of local landscape and time of year on bat-road collision risks

**DOI:** 10.1101/2020.07.15.204115

**Authors:** Charlotte Roemer, Aurélie Coulon, Thierry Disca, Yves Bas

**Author notes:** **Cite as:** Roemer C, Coulon A, Disca T, Bas Y (2020) Influence of local landscape and time of year on bat-road collision risks. bioRxiv, 2020.07.15.204115, version 3 peer-revied and recommended by Peer Community in Ecology. https://doi.org/10.1101/2020.07.15.204115. This article has been peer-reviewed and recommended by Peer Community in Ecology https://doi.org/10.24072/pci.ecology.100067.

## Abstract

Roads impact bat populations through habitat loss and collisions. High quality habitats particularly increase bat mortalities on roads, yet many questions remain concerning how local landscape features may influence bat behaviour and lead to high collision risks (e.g. influence of distance to trees, or of vegetation density). When comparing the potential danger of different road sections, the most popular method today is the use of simple bat detectors to assess the local densities of current populations at road sites. Yet, it is not known to which extent bat behaviour influences collisions (i.e. bats flying at vehicle height or on the side or above, co-occurrence of bats and vehicles). Behaviour is very rarely taken into account in practice, and this might lead to hazardous site selections for mitigation. Our goals were thus (i) to estimate how local landscape characteristics affect each of the conditional events leading to collisions (i.e. bat presence, flight in the zone at collision risk and bat-vehicle co-occurrence), and (ii) to determine which of the conditional events most contributed to collisions risks.

In this study, we recorded bat activity and characterised flight behaviour with three variables: position at collision risk, bat-vehicle co-occurrence, and flight path orientation, using acoustic flight path tracking at 66 study sites in the Mediterranean region for two to five full nights. We modelled the effect of the local landscape, i.e. in a radius of 30 m around the road (vegetation height, distance, density and orientation), road features (road width, traffic volume) and the time of year on eleven species or species groups. We built models for each conditional probability of the road collision risk (i.e. species density, presence in the zone at risk, bat-vehicle co-occurrence) and multiplied their estimates to calculate the overall collision risk.

Our results show that the local landscape had different effects on bat density and presence in the zone at collision risk. Increasing distance to trees and decreasing tree height were associated with a decrease in bat density at roads. Forests were the local landscapes where bats flew more often in the zone at collision risk. The overall collision risk was higher either in forests or at tree rows perpendicular to the road depending on species. Contrary to common preconceptions, mid-range echolocators seemed to be generally more at risk of collision than short-range or long-range echolocators. In addition, collision risk was greatest in summer or autumn for most species. Finally, bats mainly followed the road axis regardless of the type of landscape.

Our results contribute to a better understanding of bat movements in different local environments at the scale where they directly sense their surroundings with echolocation calls. Disentangling bat density from flight behaviour allowed us to better understand the temporal and spatial contributors of roadkills, and to provide guidance for road impact assessment studies.

## Introduction

Highways and main or secondary roads cover large surfaces of industrialised countries worldwide while road construction and traffic density rise continuously (Ibisch et al., 2016; van der Ree et al., 2015a). Both networks lead to troubling impacts on wildlife, namely death by collision, loss of habitat amount and quality, population fragmentation, which in turn lead to negative impacts on population survival in numerous taxa (Rytwinski and Fahrig, 2015).

To explain the direct ecological impact of roads, i.e. mortality by collision, several studies have investigated the role of road and land features on roadkill occurrence. They showed for example that road width, traffic and/or speed limit increases collisions in large mammals (Nelli et al., 2018; Neumann et al., 2012; Seiler, 2005; Valero et al., 2015), but traffic and speed limit either increased or decreased road-kills in other vertebrate taxa (Clevenger et al., 2003; D’Amico et al., 2015; Mazerolle, 2004). Studies on a variety of animal groups also found that preferred habitats for foraging or movement, described at the home-range scale (e.g. presence or absence of woodland, cropland, wetland…), are more often associated with the occurrence of road-kills (Grilo et al., 2016; Gunson et al., 2011; Malo et al., 2004).

Very few studies have investigated the effect of local habitat (i.e. within the few meters on either side of the road) on collisions. However, when mitigation measures are recommended, they often deal with the vegetation structure at this local scale (van der Ree et al., 2015b). Indeed, it is likely that the landscape in the immediate vicinity of roads affects animal movement trajectories – and, as a result, the risk of collisions. In ungulates, Meisingset et al. (2014) found that increasing road edge clearance decreased the rate of collisions. However, as the authors suggest, this is probably partly a driver effect since drivers benefitting from a better visibility will in all likelihood have more time to avoid collisions. Large animals that may be avoided by drivers represent only a very small percentage of the species impacted by collisions (D’Amico et al., 2015; Rytwinski and Fahrig, 2015) and the effects of local landscapes are likely to be species dependent, but knowledge is very scarce at the species level.

The movement of aerial animals is expected to be particularly conditioned by height, density and spatial arrangement of three-dimensional structures (Brigham et al., 1997; Norberg, 1994, 1986). For example in birds, gaps in vegetation are an important factor known to increase road collisions (Lin, 2016; Orłowski, 2008). Among aerial animals impacted by road collisions, bats are long-lived mammals with a low reproductive rate, having one offspring – exceptionally two – per year (Dietz et al., 2009). Additionally, temperate bats have suffered from an important decline of their populations in the second half of the twentieth century, which translates into a poor conservation status today (Van der Meij et al., 2015), and North-American bats have experienced dramatic declines due to white-nose syndrome, a fungal disease (Langwig et al., 2015). For these reasons, even moderate increases in mortality rates may represent a serious threat to their survival. As a result, all European bats are now under strict protection (*Council Directive 92/43/EEC on the conservation of natural habitats and of wild fauna and flora*, 1992).

Bat mortality on roads was investigated in numerous studies (Fensome and Mathews, 2016), and can locally threaten bat populations. For instance, an annual highway mortality of 5% was estimated for a colony of *Myotis sodalis* in the United States of America (Russell et al., 2009). Brinkmann et al. (2012) state that a road mortality of 3 to 7 adult females in a colony of 100 female *M. myotis* or *Rhinolophus hipposideros* could lead to a negative population growth. A good understanding of the mechanisms leading to collisions between road vehicles and bats is therefore necessary to efficiently mitigate them (Fensome and Mathews, 2016).

At the home range scale, several studies showed that collisions involving bats are concentrated in habitats classified as favourable for foraging and commuting (e.g. water bodies, forests and riparian habitats) (Gaisler et al., 2009; Lesiński, 2007; Medinas et al., 2019, 2013). At the local scale, it is suspected that the orientation of tree lines and vegetation structure (i.e. height, density and distance from road edge) direct bat movement, and are consequently major factors of road collision risks (Fensome and Mathews, 2016). The influence of vegetation structure on bat activity has been relatively well studied in the literature (Kelm et al., 2014; Pourshoushtari et al., 2018; Toffoli, 2016; Verboom and Spoelstra, 1999). However, the influence of local landscape on bat movement has been almost exclusively described from visual observations (Arthur and Lemaire, 2015; Dietz et al., 2009). Some studies on roads suggest that increasing distance to surrounding trees decreases bat road crossing frequency, and that increasing tree height elevates bat crossing height (hence reducing the risk of collisions with vehicles) (Abbott, 2012; Bennett and Zurcher, 2013; Russell et al., 2009). But small sample sizes and poor taxonomic resolution limit the generalisation of these results. Moreover, Bennett and Zurcher (2013) considered bat trajectories initially directed perpendicular to the road and determined that vegetation structure and vehicle presence influenced bat decisions to cross the road or to fly away. But they did not take into account bats flying parallel to the road axis, although this behaviour may be a determinant factor of collisions, because bats flying parallel to the road axis may fly at risk of collisions for dozens of meters, while crossing a road only implies flying at risk of collision for a few meters. Mitigation measures to reduce collisions are also mainly designed for bats crossing roads (Elmeros et al., 2016), although it is not known to which extent bats may follow the road axis or cross roads, depending on the habitat context.

Road collision risks in a species depend on (1) its local density, (2) the proportion of time spent in the zone at collision risk and (3) the simultaneous presence of bats and vehicles in the zone at collision risk (Jaeger et al., 2005; Zimmermann Teixeira et al., 2017). It is therefore necessary to take each of these conditional events into account when investigating road collisions. Indeed, when comparing two different road locations within, different landscape features, a higher bat acoustic activity (used as a proxy of bat density) does not necessarily lead to a higher proportion of flights at collision risk for all species (see Abbott et al., 2012). In addition, even if more individuals are at risk of collision (or if mortality is higher) at one site compared to another, this does not necessarily mean that this site should be selected for mitigation. Indeed, local populations can be dramatically reduced due to road mortality year after year, and a measure of per capita mortality risk is essential to correctly identify dangerous locations and avoid wrong recommendations for the siting of mitigation measures (Zimmermann Teixeira et al., 2017). Per capita mortality is also a very useful tool to prioritise conservation actions in function of the susceptibility of species to anthropogenic impacts. For instance, bats of the *Nyctalus* genus are particularly susceptible to wind turbine collisions because a high proportion of the individuals are victims of collisions (Roemer et al., 2017); to spare their populations, wind energy planning should therefore avoid areas where these species are extant.

The aim of our study was to assess the effects of the local habitat, coupled with bat density and movement patterns, on road collision risks. In order to provide species-specific answers, our analyses were mostly performed at the species level, using the guild level only for species with small sample sizes. In addition, one of our goals was to provide a proxy for bat guilds susceptibility to road collisions independently of their population sizes. We used acoustic monitoring to detect bat passes and car passes, and acoustic flight path tracking to locate bat echolocation calls in three dimensions. This method allows reconstructing three-dimensional flight paths, and then model separately: (1) bat species density, (2) a probability of flight at collision risk, (3) a probability of bat-vehicle co-occurrence and (4) a probability to fly parallel or perpendicular to the road.

We expected bat density to be the main factor influencing collision risks in some contexts (for example in habitats classified as favourable for bat foraging and commuting such as forests and riparian habitats) (Gaisler et al., 2009; Lesiński, 2007; Medinas et al., 2019, 2013), but we expected the proportion of individuals flying in the zone at collision risks to be the main factor in other contexts (especially in forested areas and when vegetation grows closer to the road, acting as a conduit (Kalcounis-Rueppell et al., 2013) and possibly forcing bats to fly over the road). In addition, we expected a correlation between the orientation of bat trajectories and the orientation of linear vegetation (Holderied, 2006; Kalcounis-Rueppell et al., 2013; Limpens and Kapteyn, 1991) and a larger proportion of individuals flying in the zone at collision risk for short-range echolocators than for mid-range echolocators and long-range echolocators, reflecting the vertical niches of those species (Roemer et al., 2019).

## Material and Methods

### Study sites and description of the local landscape

The study took place in 2016 and 2017 in the French Mediterranean lowland region. This area is composed of a mosaic of garrigues, cultivated areas (often vines), young forests of oaks and pines, and urban areas consisting of small traditional villages and large cities with extensive conurbation. The national and departmental road network is built around one main highway going from the east to the west and following, the French southern coastline (**Figure 1**). Four other highways link this main highway to the inner lands through large natural valleys.

**Figure 1:**
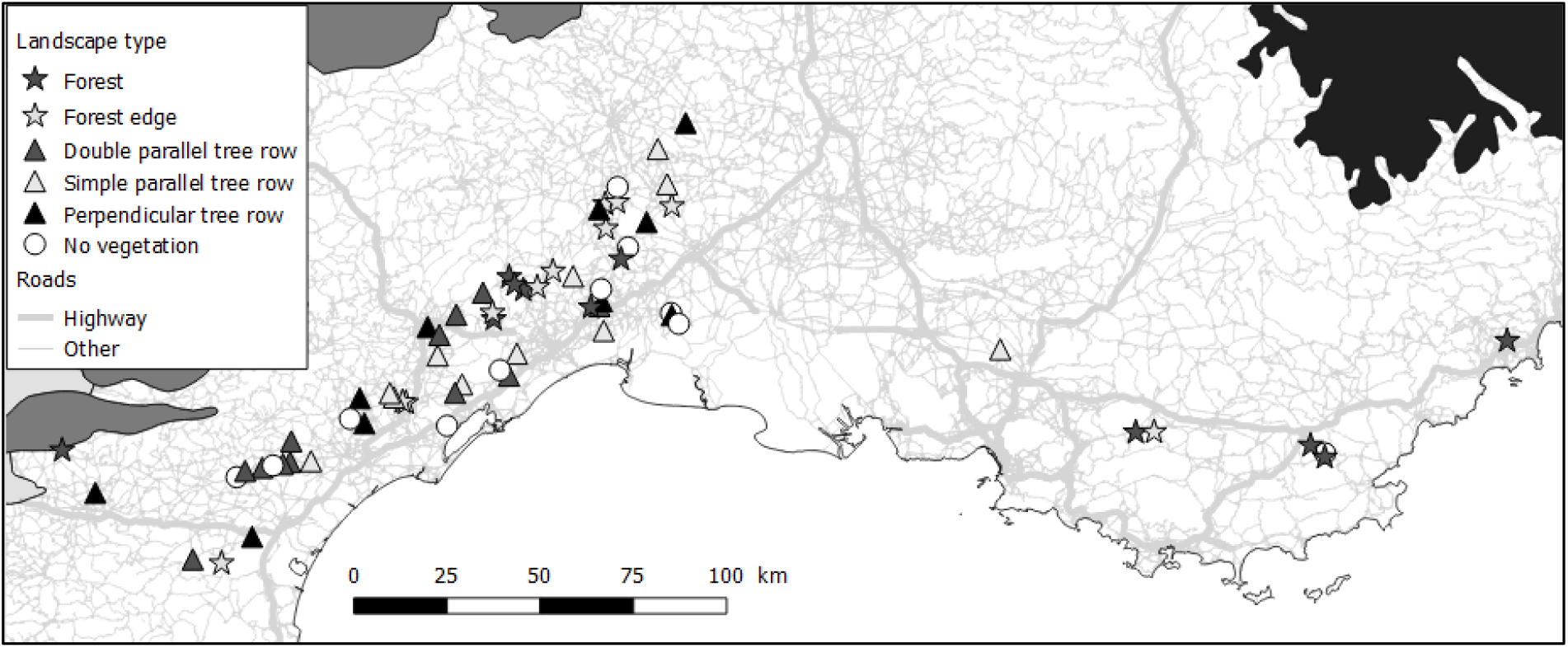
Geographical distribution of the 66 sampling sites in the secondary road network of the French Mediterranean lowland region. The symbols of the sampling sites represent the type of local landscape they belong to. Biogeographical regions are filled in colour: white: Mediterranean; grey: Continental; black: Alpine. Road network source: ROUTE500 from the Institut National de l’Information Géographique et Forestière (2017).

Bat behaviour was recorded at 66 sites (**Figure 1**, supplementary table 1) at national or departmental roads, for a minimum of two nights per site, but recordings could continue up to five nights per site (mean = 2.6 nights +/− SD 0.9) depending on the schedule of the field worker and on battery strength. Sampling took place between the beginning of May and mid-October, depending on the study site. The local landscape was described within a radius of 30 m, equivalent to the sonar range of mid-range echolocating bats (Holderied and von Helversen, 2003). This scale was chosen under the assumption that bats adapt their flight movements according to the environment perceived acoustically. At most study sites, landscape description would have been similar with a 100 m radius, which corresponds to the sonar range of long-range echolocators (Holderied and von Helversen, 2003).

Study sites were chosen so as to reach a balanced representation of six major types of road landscapes in the study area: simple parallel tree rows, double parallel tree rows, perpendicular tree rows, forests, forest edges and no vegetation taller than 1.5 m (Table 1, Figure 1, **Figure A 1**). Tree species were very often associated with a type of landscape: 70 % of simple and double parallel tree rows were plane trees (*Platanus sp*.), and more rarely olive trees (*Olea sp.*), Celtis (*Celtis sp*.), Aleppo pines (*Pinus halepensis*) or mulberries (*Morus sp.*). Forests and forest edges consisted in 80 % oaks (*Quercus ilex*, *Q. pubescens*, *Q. coccifera* and/or *Q. suber*), and 20 % Aleppo pines (*P. halepensis*). Perpendicular tree rows were a mix of Mediterranean riparian species (mostly *Fraxinus* sp., *Populus* sp., *Quercus* sp. and/or *Arundo donax*) typically associated with temporary watercourses. The category “no vegetation” consisted in land either occupied by vines, wheat, recently ploughed or left uncultivated. Pastureland was almost non-existent. Parallel tree rows had gaps of about 10 to 20 m between trees while the other types of vegetated landscapes had little or no gaps.

**Table 1:** Description of variables used to model bat density, bat position at collision risk, and flight path orientation.

| *Variable* | *Description* |
| --- | --- |
| **Landscape type** | **Simple parallel tree row**<br>One row of trees arranged linearly and parallel to the road. |
|  | **Double parallel tree row**<br>Two rows of trees arranged linearly at each side and parallel to the road. |
|  | **Perpendicular tree row**<br>One row of trees at each side and perpendicular to the road. Associated with small seasonal streams where water was either absent at the time of monitoring, or likely not accessible for bats due to dense tree cover. |
|  | **Forest**<br>Dense tree patch at each side of the road. |
|  | **Forest edge**<br>Dense tree patch at one side of the road. |
|  | **No vegetation**<br>No vegetation taller than 1.5 m at each side of the road. |
| **Road width** | Distance between both outer edge lines of the roadway. |
| **Traffic volume** | Mean number of vehicles per night. |
| **Distance to tree foliage** | Mean distance between road outer edge line and tree foliage over all present trees. If foliage runs over the road, distance is negative. |
| **Tree height** | Mean tree height from ground to canopy over all present trees. |

All sites were situated in lowlands, on two-lane asphalt roads of 4 to 8 m wide, on straight portions (at least 200 m without curvature on each side of the sampling point), where vehicles were allowed to drive up to 90 km/h. Several features were avoided: (1) artificial street lights and urban areas (the smallest distance to lit streets and urban areas was 300 m), (2) important three-dimensional structures, such as electric poles, (3) highways (the smallest distance to a highway was 1.1 km), (4) water bodies or wetlands other than the small streams sampled in the category “perpendicular tree rows” (the shortest distance to water was 100 m) and (5) sparse trees within the landscape matrix. The minimum distance between study sites was 500 m. Monitoring was performed exclusively during nights with optimal weather conditions for bat activity (temperature: mean = 20.6 +/− 6.5 °C, min = 8 °C, max = 34.9 °C; wind speed: mean = 7.5 +/− 8 km/h, min = 0 km/h, max = 31 km/h; accumulated rain per night: mean = 0.2 +/− 1.3 mm, min = 0 mm, max = 11 mm). However, the percentage of visible moon (mean = 49.2 +/− 35.8 %, min = 1 %, max = 99 %) was not a criterion we could control because of the time constrained field work schedule.

Four secondary landscape characteristics likely to affect flight behaviour were measured at each study site: road width, traffic volume, distance between road and tree foliage, and tree height (**Table 1**). Measurements were made with a laser telemeter. Traffic volume was calculated using the TADARIDA-L software (Bas et al., 2017) to identify and count vehicle passes. Sound event detection was done in the low frequency mode, and any acoustic sequence of 5 s or less that contained an uninterrupted sound event with a duration superior to 1.2 s was counted as one vehicle pass, even if several vehicles followed each other very closely. This threshold was chosen based on a verification of false and true positives and negatives on 100 random sound sequences from different study sites stratified by sound duration (unpublished data).

### Bat acoustic monitoring

On each site, two pairs of microphones (either SMX-US or SMX-U1 (Wildlife Acoustics, USA), or BMX-US (Biotope, France)) were plugged into two SM2BATs or SM3BATs (Wildlife Acoustics, USA), each connected to a GPS unit used to timely synchronise recorders. Microphones were either mounted on wooden poles (at a maximum height of 4 m) or attached to vegetation (at a minimum height of 20 cm) (**Figure A 1**). Microphone pairs were installed on each side of the road (0.5 – 4 m distance from the road edge) in arbitrarily shaped non-coplanar microphone arrays. Depending on the study site, minimum distance between microphones was 5.1 m and maximum distance was 22.6 m. Recorders were programmed to start each day 30 min before sunset and to stop 30 min after sunrise. Gain was set at 36 dB for SMX-US and BMX-US microphones, or at 0 dB for SMX-U1 microphones. Sampling rate was set at 192 kHz, trigger at 6 dB above background noise and trigger window at 2.5 sec.

Species identification was performed based on echolocation calls, which carry enough information to allow the identification of the majority of European bat species, depending on the quality and the context of the recordings (Barataud, 2015). We used the SonoChiro software (Biotope/MNHN, France) to automatically sort sequences by species, and then verified most of the sequences manually on Syrinx (John Burt, USA) (except for sequences classified as *Pipistrellus* which are too numerous for a detailed verification, and because SonoChiro has a very low error rate for *P. pipistrellus* and *P. pygmaeus* in the Mediterranean region according to our experience). *Plecotus* species were grouped in *Plecotus* sp., *Myotis blythii* and *M. myotis* were grouped in *M. blythii/myotis*, and *Pipistrellus kuhlii* and *P. nathusii* were grouped in *Pipistrellus kuhlii/nathusii*. Acoustic sequences that could not be identified to the species level or groups were left unidentified (0.15 % of all bat passes). From our knowledge of bat assemblages of France Mediterranean lowlands (unpublished mist-netting data), we expect the last group to contain a very large majority of *P. kuhlii*, and the *Plecotus* group to contain a very large majority of *P. austriacus*.

### Three-dimensional positioning of bat calls

Bat three-dimensional flight paths were generated from the three-dimensional source location of echolocation calls recorded on the four microphones. After species acoustic identification, call location was achieved by (1) measuring time of arrival differences (TOAD) of each call between pairs of microphones and deducing the coordinates of the sound source by comparing those field TOAD (TOAD_F_) with theoretical TOAD (TOAD_T_). Indeed, since the speed of sound in the air is known (here we approximated it to 340 m/s), TOADs of a sound source recorded by at least four microphones can be used to calculate the location of the source (see Koblitz, 2018).

TOAD_F_ were calculated by measuring the starting time of bat calls using the SonoChiro software (Biotope/MNHN, France). Call association between pairs of microphones was achieved using the R (R Core Team, 2014) function find.matches of the Hmisc package (Harrell, 2018). Because there are four microphones, six TOAD_F_ per call are calculated. TOAD_T_ were calculated for each simulated point of a matrix of 40 × 40 × 40 m with a one-meter resolution and centred around the centroid of the 4 microphones, inputting the same microphone configurations as the ones used in the field. The dimensions of this matrix were chosen according to the spatial range of our equipment (i.e. maximal distance of detection of a bat position) for the location of middle-range echolocators (e.g. *Pipistrellus pipistrellus*). This range is dependent on the acoustic range of the, individual recorded and the position of the individual in relation to the microphones, i.e. accuracy is maximal at the centre of the device and minimal at the far edges.

The position of the bat was deduced from the comparison of the differences between the six TOAD_F_ and the six TOAD_T_ using the R (R Core Team, 2014) function find.matches of the Hmisc package (Harrell, 2018). The closest TOAD match was selected as a candidate bat position. During test calibrations of our setting with different microphone configurations, we calculated that TOAD_F_ resulting in a position more than 10 m away from the centroid of the microphones had a difference with the real position larger than one meter. Imprecise positions were systematically reconstructed away from the centroid, which means for example that a bat flying in reality at 15 m from the centroid could be located with our device at 17 m (away from the centroid), but never at 13 m (toward the centroid). We therefore rejected any field position found at more than 10 m from the centroid of the microphones and did not use them for further analyses.

### Grouping of calls in individual flight trajectories

Calls were then attributed a flight trajectory ID using successive filters. During the first round, a same temporary ID was first given to all calls separated by less than two seconds and the flight speed between the preceding and the actual call was calculated in the X and Y dimensions. Several rounds were then run successively. At each round, to keep their temporary ID, calls had to (1) have a peak frequency differing by less than 5 kHz from the median peak frequency of all calls within the same ID (2) be separated by less than 2 seconds from the preceding call (3) be preceded and followed by positions conferring a speed lower than 20 m/s (i.e. the maximum possible speed of flying bats (Holderied and Jones, 2009; Popa-Lisseanu, 2007). Otherwise, calls were attributed a new (unique) temporary ID and went through a new round of filtering. Successive rounds were applied until all IDs were stabilized. Flight trajectories with less than three calls were not considered as a full flight path and were classified as non-located bat passes (**Figure 2**). R scripts and tables are available at https://github.com/Charlotte-Roemer/bat-road-collision-risks.

**Figure 2:**
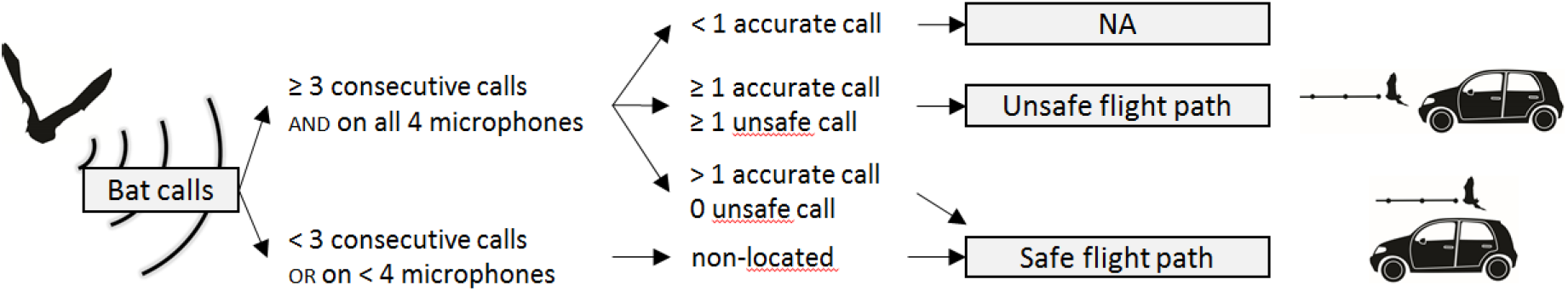
Process of classification of flight trajectory at collision risk. NA: non-available data. The area where calls are accurate is 10 m around the centroid of microphones. Unsafe call: located at vehicle height and above the road.

### Definition of collision risk and calculation of flight path orientation

Each successfully located bat position above the road and at vehicle height (< 5 m) (Berthinussen and Altringham, 2012) was classified as ‘unsafe’ (**Figure 2**). All other successfully located positions were classified as ‘safe’. Bat calls which were not recorded by all four microphones at once – and that could therefore not be precisely located – were assumed to be far from the microphones’ centroid and thus probably far from the road and hence also classified as ‘safe’. For the same reason, positions potentially not successfully located (> 10 m from microphones centroid) were disregarded to avoid location errors. Since this error rate is similar across landscape types, we do not expect any resulting bias. If any of the bat positions within a flight trajectory was unsafe, the complete flight trajectory was classified as unsafe, otherwise it was classified as safe. Flight trajectories with less than three calls were assumed to be far from the microphones’ centroid and thus probably far from the road and hence also classified as ‘safe’ (**Figure 2**).

We then calculated the angle between the road axis and the axis of the vector linking the first to the last position of each flight paths. Trajectories were classified in two categories: 0-45° = parallel; 45-90° = perpendicular to the road.

### Response variables

We tested how the local landscape affects the different determinants of collision risks, building one model for each of them: (1) local species density (2) the proportion of flights in the zone at risk (3) bat-vehicle co-occurrence and (4) flight path orientations (Figure 3). To summarise the results, the estimates of the first three models (i.e. quantitative models) were multiplied to obtain as a product (5) the number of bat passes at collision risk per night, depending on the characteristics of the local landscape.

**Figure 3:**
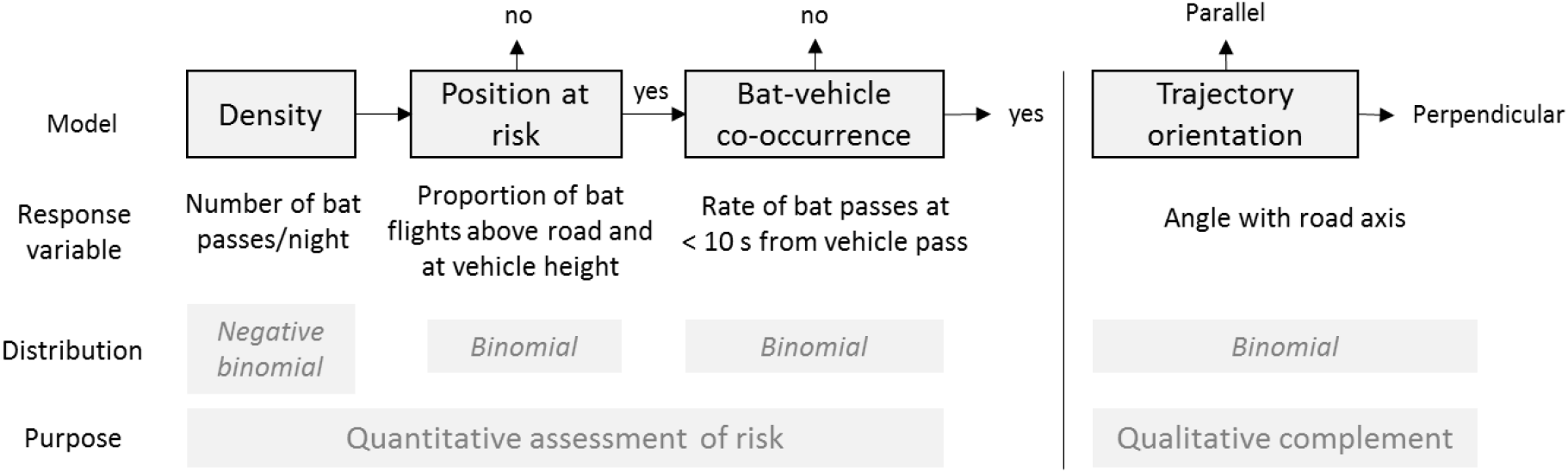
Successive steps in modelling of bat density and flight behaviour in function of landscape variables. If the flight path is at risk, position at risk = yes. If the time lag between the bat pass and the vehicle pass is inferior to 10 s, bat-vehicle co-occurrence = yes.

#### (1) Local bat density

The density of the most common species in our dataset (i.e. occurrence per night larger than 30 % and occurrence per site larger than 50 %) was modelled as a negative binomial distribution. The response variable was the median number (among the four microphones) of five second intervals per night within which a species was identified. This acoustic activity was then used as a proxy of bat density (number of acoustic sequences per night within the acoustic range of the setting) (Froidevaux et al., 2017).

#### (2) Probability of bats flying through the zone at risk

The probability of trajectories to be in the zone at risk (i.e. at vehicle height and above the road) was modelled using the risk status of each trajectory as the binomial response variable (0 = safe; 1 = unsafe) (**Figure 2**).

#### (3) Bat-vehicle co-occurrence

The probability of bats flying through the zone at risk gives a spatial evaluation of risk. To make a more precise risk assessment, bat-vehicle co-occurrence (i.e. temporal evaluation of risk) was also modelled. For bat flight trajectories at risk only, a proxy for the probability of bats avoiding vehicles was modelled using bat-vehicle co-occurrence as the binomial response variable (1 = bat-vehicle co-occurrence; 0 = presence of a bat while absence of vehicle). To do this, the time lag between an acoustic sequence containing a bat and the closest sequence containing a vehicle pass was calculated using the function find.matches of the Hmisc package (Harrell, 2018). If the time lag was lower than 10 s, we considered that there was a bat-vehicle co-occurrence (1). If the time lag was higher than 10 s, we considered that a bat was present during the absence of a vehicle (0).

#### (4) Flight path orientation

For all bat flight trajectories, the proportion of flight paths parallel to the road axis was modelled using flight orientation as a binomial response variable (0 = perpendicular; 1 = parallel). This model is not a quantitative estimation of the collision risk at a road section, since the road sections that we studied were approximatively squared, and thus the orientation of bat trajectories does not influence the time spent at risk of collision. This model was therefore made to provide a qualitative estimation of the collision risk that can help the design of mitigation measures. Even if bats flying parallel and above the road do fly for a longer among of time at risk of collision than bats crossing roads, in the case of our study, we assessed collision risks relatively to a road section and not relatively to a bat individual.

#### (5) Number of bat passes at collision risk per night

If an explanatory variable was selected in several models, then each of those models gives a partial evaluation of bat collision risks on roads. In fact, all quantitative models succeeding the density model can be interpreted as conditional probabilities that an individual is at risk of collision. Thus, the number of bat passes at risk of collision on a road section can be computed by the multiplication of all outputs of the quantitative models:

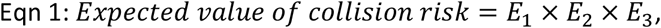

where:

E_1_ = Prediction of the number of bats present on site per night

E_2_ = Prediction of the probability that a detected bat flies in the volume at collision risk

E_3_ = Prediction of the probability that a bat in the volume at risk co-occurs with a vehicle pass

To estimate the confidence intervals of this product, we needed a large number of responses for each value along the gradient of each predictor for each response variable. For this, we first simulated responses according to model estimates and their standard error using the rnorm function (R Core Team, 2014): 20,000 replicates for each of 60 values along a gradient for a given predictor. For a given predictor, if one E_1_, E_2_ or E_3_, was missing, meaning that the predictor was not selected in one of the models, we used the mean value of the original observations instead. If two or all three of E_1_, E_2_ or E_3_ were missing for a given predictor, we did not compute their product.

Since our results apply for road sections 20 m in length, we multiplied the expected mean number of bat passes at collision risk by 50 to obtain a mean number of bat passes at risk of collision per kilometre and per night. To compare bat guilds susceptibility to road collisions, we multiplied *E*_2_ × *E*_3_ this result is an index of susceptibility to road collisions that is independent of local population densities.

### Model selection

We used the R (R Core Team, 2014) package glmmTMB (Brooks et al., 2017) to model each response variable in generalised linear mixed models (GLMM). When sample size of a given species was too small, we did not model the species response. In addition, three bat guilds were created based on the adaptation of species to clutter of the environment, which is strongly linked to sonar features (Aldridge and Rautenbach, 1987; Denzinger et al., 2018). Species were thereby split into the guilds “short-range echolocator” (SRE), “mid-range echolocator” (MRE) or “long-range echolocator” (LRE) according to the definition of Frey-Ehrenbold et al. (2013) (see **Table A 1** for complete list).

All descriptive variables were normalised if necessary and scaled to follow a normal distribution and to compare their effects. Thus, distance to vegetation and traffic were normalised using the square root function. Variables considered for fixed effects were landscape type, road width, traffic volume, distance to tree foliage and tree height (**Table 1**). We first calculated the correlation coefficients between predictors using the corrplot function of the stats package in the R program (R Core Team, 2014). Tree height and distance to tree foliage were correlated (r = −0.57), as well as road width and traffic volume (r = 0.64). We excluded road width for further analysis and created the possibility to select either tree height or distance to tree foliage (but not both) during stepwise model selection (see next paragraph). Candidate predictors were also included in simple interactions with each other. In addition, Julian day was included as a fixed quadratic effect to account for seasonal variations in bat density and flight behaviour. Site ID was introduced as a random effect.

An upward stepwise model selection was performed to select the relevant variables (except for Julian day which was part of the null model). We operated an upward model selection because the full model led to overfitting for species with a low occurrence. At each step of model selection, the VIF (Variance inflation factor), which quantifies the degree of multicollinearity in least square regression analyses, was calculated. If any of the selected variables had a VIF > 3 (Heiberger and Holland, 2004; Zuur et al., 2010), the model was not considered as a candidate model. At each step of model selection, the model with the smallest Akaike’s information criterion for small sample sizes (AICc) was considered. This model was retained and selected if its AICc was at least inferior by two points to the AICc of the best model of the previous step (supporting that the newly added parameter is informative) (Arnold, 2010).

For each retained model, we checked the uniformity of the residuals using the DHARMa package (Hartig, 2018). Goodness of fit, autocorrelation, overdispersion and zero-inflation (for density data only) were checked and revealed no problematic situation.

## Results

In total, 122,294 bat passes were recorded and identified at the group or species level, from which 30,954 successful flight trajectories could be located (**Table A 1**). Because of technical problems on two study sites (one of the two recorders was once destroyed by a rotary flail and once displaced by someone), flight path tracking could not be carried out and these sites were used for modelling bat density only. The density of nine species and three species groups (*Pipistrellus kuhlii/nathusii*, *Plecotus sp*. and *Myotis blythii/myotis*) could be modelled, but their flight behaviour (i.e. presence at risk, bat-vehicle co-occurrence and flight path orientation) could not be modelled for all of the species or species groups, due to the lack of data.

Models showed no convergence problems during selection, except for *E. serotinus* (model position in the zone at risk, for interactions), *M. daubentonii* (model bat-vehicle co-occurrence, for landscape type), H. savii (model trajectory orientation, for landscape type) and *M.myotis/blythii* (model bat-vehicle co-occurrence, for landscape type; model trajectory orientation, for landscape type). When model convergence failed, the model could not be built and was not considered for selection.

### Model 1 – Bat density

Landscape type had an important influence on bat density (**Table 2**). It was selected in the model of four species. Density was much higher at perpendicular tree rows for *Pipistrellus* species and was higher at forested landscapes for *H. savii* (**Figure 4**). Increasing distances to tree foliage were associated with a decrease in bat density for five species (*M. daubentonii, P. pipistrellus*, *P. pygmaeus*, *M. schreibersii* and *N. leisleri*) and for the MRE guild, while it was associated with an increase for *Plecotus* species (**Figure 5**). Increasing tree height was associated with an increase in the density of *M. blythii/myotis* and of the LRE guild (**Table 2**). With an increasing traffic volume, the density of *Plecotus sp*. and of the SRE guild decreased (**Table 2**). Throughout the year, species density showed a typical peak in mid-summer, except for *P. pygmaeus*, *M. schreibersii, Plecotus* sp. and *N. leisleri*, that were more active in the autumn (**Figure A 2**).

**Table 2:** Summarised statistical results of the negative binomial distributed GLMM for the density of each species. 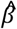 = estimate. SE = standard error. p = significance of p value. Species names are given with the three first letters of the species and genera. Dist.tree = distance to tree foliage. Tree.H = tree height. DPT = double parallel tree rows. F = forest. FE = forest edge. PT = perpendicular tree rows. SPT = simple parallel tree rows. Intercept is for NV (no vegetation) landscape type. LRE: long-range echolocators. MRE: mid-range echolocators. SRE: short-range, echolocators. There is a total of 163 nights of observations.

| Variable |  | Eptser |  |  | Hypsav |  |  | Minsch |  |  | Myodau |  |  | Myoema |  |  | Myobly/myo |  |  | Nyclei |  |  | Pipkuh/nat |  |  | Pippip |  |  | Pippyg |  |  | Plesp |  |  |
| --- | --- | --- | --- | --- | --- | --- | --- | --- | --- | --- | --- | --- | --- | --- | --- | --- | --- | --- | --- | --- | --- | --- | --- | --- | --- | --- | --- | --- | --- | --- | --- | --- | --- | --- |
| Type | | $\beta$ | SE | p | $\beta$ | SE | p | $\beta$ | SE | p | $\beta$ | SE | p | $\beta$ | SE | p | $\beta$ | SE | p | $\beta$ | SE | p | $\beta$ | SE | p | $\beta$ | SE | p | $\beta$ | SE | p | | | |
| Intercept |  | 0.64 | 0.48 |  | 2.64 | 0.50 | *** | -0.17 | 0.56 |  | -0.76 | 0.41 | . | 0.00 | 0.28 |  | 0.32 | 0.39 |  | 2.24 | 0.22 | *** | 5.50 | 0.19 | *** | 4.67 | 0.29 | *** | 3.03 | 0.44 | *** | 0.26 | 0.23 |  |
| Dist.tree |  |  |  |  |  |  |  | -0.58 | 0.23 | * | -0.92 | 0.29 | ** |  |  |  |  |  |  | -0.91 | 0.17 | *** |  |  |  | -0.29 | 0.13 | * | -0.80 | 0.20 | *** | 0.42 | 0.16 | * |
| Tree.H |  |  |  |  |  |  |  |  |  |  |  |  |  |  |  |  | 1.21 | 0.33 | *** |  |  |  |  |  |  |  |  |  |  |  |  |  |  |  |
| Traffic |  |  |  |  |  |  |  |  |  |  |  |  |  |  |  |  |  |  |  |  |  |  |  |  |  |  |  |  |  |  | -0.44 | 0.17 | * |  |
| Landscape type | DPT |  |  |  | -0.20 | 0.62 |  | 1.04 | 0.72 |  |  |  |  |  |  |  |  |  |  |  |  |  |  |  | 0.44 | 0.39 |  | 0.62 | 0.59 |  |  |  |  |  |
|  | F |  |  |  | 0.63 | 0.64 |  | 3.34 | 0.71 | *** |  |  |  |  |  |  |  |  |  |  |  |  |  |  | 0.82 | 0.38 | * | 0.06 | 0.58 |  |  |  |  |  |
|  | FE |  |  |  | 0.76 | 0.63 |  | 3.34 | 0.72 | *** |  |  |  |  |  |  |  |  |  |  |  |  |  |  | 0.67 | 0.39 | . | 0.14 | 0.59 |  |  |  |  |  |
|  | PT |  |  |  | -1.42 | 0.67 | * | 3.26 | 0.72 | *** |  |  |  |  |  |  |  |  |  |  |  |  |  |  | 2.22 | 0.39 | *** | 1.87 | 0.59 | ** |  |  |  |  |
|  | SPT |  |  |  | -1.35 | 0.67 | * | 2.15 | 0.75 | ** |  |  |  |  |  |  |  |  |  |  |  |  |  |  | -0.16 | 0.40 |  | -0.11 | 0.61 |  |  |  |  |  |
| Julian Day |  | -0.19 | 0.30 |  | -0.21 | 0.18 |  | 0.42 | 0.15 | ** | 1.17 | 0.32 | *** | -0.30 | 0.18 |  | -0.85 | 0.28 | ** | 0.40 | 0.14 | ** | -0.09 | 0.13 |  | 0.40 | 0.10 | *** | 0.74 | 0.14 | *** | 0.53 | 0.15 | *** |
| Julian Day<sup>2</sup> |  | -0.96 | 0.30 | ** | -0.99 | 0.19 | *** | 0.36 | 0.14 | ** | -0.58 | 0.26 | * | -0.44 | 0.19 | * | -0.75 | 0.23 | ** | 0.28 | 0.14 | * | -0.50 | 0.12 | *** | -0.58 | 0.09 | *** | 0.07 | 0.13 |  | 0.41 | 0.14 | ** |

Table 2 (continued)
| Variable |  | SRE |  |  | MRE |  |  | LRE |  |  |
| --- | --- | --- | --- | --- | --- | --- | --- | --- | --- | --- |
| Type | | $\beta$ | SE | p | $\beta$ | SE | p | $\beta$ | SE | p |
| Intercept |  | 3.09 | 0.16 | *** | 5.56 | 0.26 | *** | 3.17 | 0.19 | *** |
| Dist.tree |  |  |  |  | -0.31 | 0.12 | ** |  |  |  |
| Tree.H |  |  |  |  |  |  |  | 0.71 | 0.15 | *** |
| Traffic |  | -0.36 | 0.14 | ** |  |  |  |  |  |  |
| Landscape type | DPT |  |  |  | 0.62 | 0.35 | . |  |  |  |
|  | F |  |  |  | 1.35 | 0.34 | *** |  |  |  |
|  | FE |  |  |  | 1.01 | 0.35 | ** |  |  |  |
|  | PT |  |  |  | 1.72 | 0.35 | *** |  |  |  |
|  | SPT |  |  |  | 0.53 | 0.36 |  |  |  |  |
| Julian Day |  | 0.19 | 0.10 | . | 0.18 | 0.09 | * | 0.30 | 0.13 | * |
| Julian Day<sup>2</sup> |  | -0.29 | 0.10 | ** | -0.30 | 0.08 | *** | -0.05 | 0.12 |  |

P < 0.1 = . P < 0.05 = \* P < 0.01 = \*\* P < 0.001 = \*\*\*

**Figure 4:**
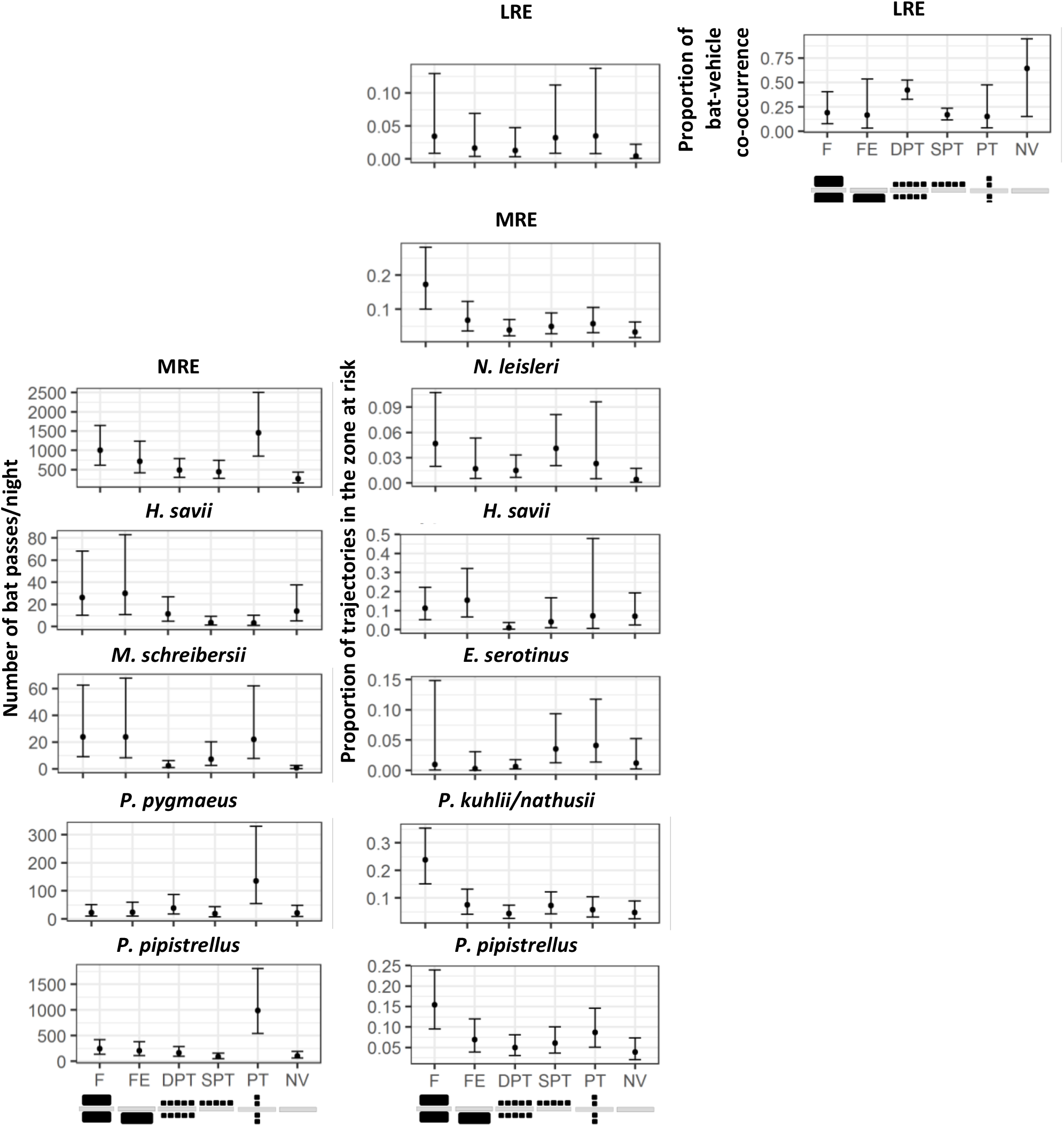
Predicted effects of landscape type on density (left), proportion of trajectories at risk (middle) and predicted proportion of bat-vehicle co-occurrence (trajectories at risk and at less than 10 s from a vehicle pass) (right). 95% confidence intervals are shown. Only the effects present in the final models are shown. Bottom figures represent landscape type viewed from the top (road in light grey and trees in black). LRE: long-range echolocators. MRE: mid-range echolocators. F = forest. FE = forest Edge. DPT = double parallel tree rows. SPT = simple parallel tree rows. PT = perpendicular tree rows. NV = no vegetation.

**Figure 5:**
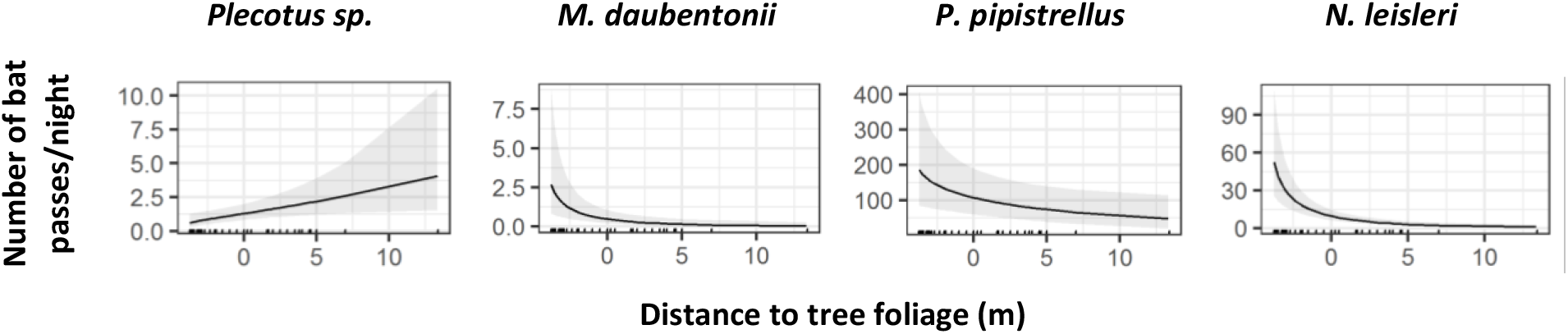
Predicted effect of distance to foliage on bat density for a selection of four species. 95% confidence intervals are shown. Ticks in x axis represent sampled values. Negative values mean that foliage was running over the road.

### Model 2 – Bat presence in the zone at risk

Landscape type also greatly influenced the proportion of bat positions in the zone at risk. It was selected in five of the ten species-specific models, and in two of the guild models. The proportion of positions at risk was generally higher in forests and lower without trees (**Figure 4** and **Table 3**).

**Table 3:**
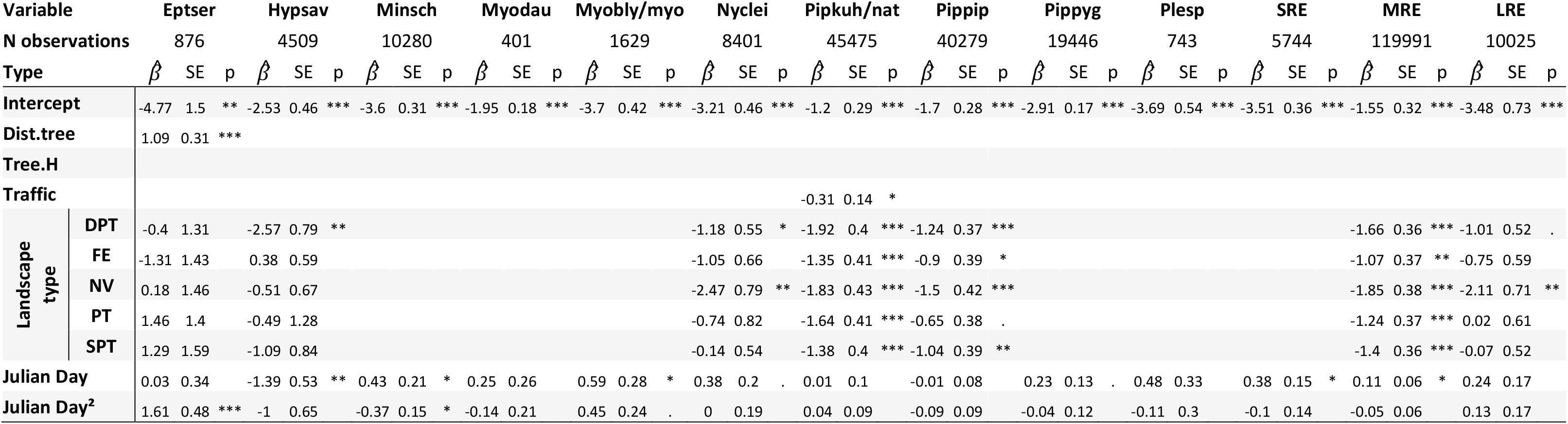
Summarised statistical results of the binomial distributed GLMM for the proportion of trajectories in the zone at risk for each species. 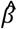 = estimate. SE = standard error. p = significance of p value. Species names are given with the three first letters of the species and genera. Dist.tree = distance to tree foliage. Tree.H = tree height. DPT = double parallel tree rows. NV = no vegetation. FE = forest Edge. PT = perpendicular tree rows. SPT = simple parallel tree rows. Intercept is for F (forest) landscape type. LRE: long-range echolocators. MRE: mid-range echolocators. SRE: short-range echolocators.

An increasing distance to tree foliage was associated with an increase in the presence at risk for *E. serotinus* (**Table 3**). Increasing traffic density was associated with a decrease in the proportion of flights in the zone at risk for *P. kuhlii/nathusii* (**Table 3**). Throughout the year, the different species displayed quite different patterns in presence at risk, but all three guilds showed a tendency for a higher proportion of flights in the zone at risk toward the end of the year (**Figure A 2**).

### Model 3 – Bat-vehicle co-occurrence for trajectories in the zone at risk

Landscape type was only selected in the model for the LRE (**Table 4**). For this guild, double parallel tree rows were associated with higher bat-vehicle co-occurrence than simple parallel tree rows. Increasing tree height was associated with an increase in bat-vehicle co-occurrence in *M. daubentonii, P. pipistrellus, P. pygmaeus* and the MRE guild (**Table 4**). An increasing traffic density was associated to an increase in bat-vehicle co-occurrence for all *Pipistrellus* species and for *M. myotis/blythii* (**Table 4**). The MRE guild had a higher rate of bat-vehicle co-occurrence than the SRE guild (**Figure 6**). Season had different effects on bat-vehicle co-occurrence according to species and guilds (**Table 4**, **Figure A 2**).

**Table 4:** Summarised statistical results of the binomial distributed generalised linear mixed effect models (GLMM) for bat-vehicle co-occurrence for each species. 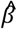 = estimate. SE = standard error. p = significance of p value. Species names are given with the three first letters of the species and genera. Dist.tree = distance to tree foliage. Tree.H = tree height. DPT = double parallel tree rows. NV = no vegetation. FE = forest Edge. PT = perpendicular tree rows. SPT = simple parallel tree rows. Intercept is for F (Forest) landscape type. LRE: long-range echolocators. MRE: mid-range echolocators. SRE: short-range echolocators.

| Variable |  | Eptser |  |  | Hypsav |  |  | Minsch |  |  | Myodau |  |  | Myobly/myo |  |  | Nyclei |  |  | Pipkuh/nat |  |  | Pippip |  |  | Pippyg |  |  | SRE |  |  | MRE |  |  | LRE |  |  |
| --- | --- | --- | --- | --- | --- | --- | --- | --- | --- | --- | --- | --- | --- | --- | --- | --- | --- | --- | --- | --- | --- | --- | --- | --- | --- | --- | --- | --- | --- | --- | --- | --- | --- | --- | --- | --- | --- |
| N observations |  | 35 |  |  | 869 |  |  | 661 |  |  | 46 |  |  | 60 |  |  | 432 |  |  | 5155 |  |  | 5291 |  |  | 1356 |  |  | 213 |  |  | 13332 |  |  | 480 |  |  |
| Type | | $\hat{\beta}$ | SE | p | $\hat{\beta}$ | SE | p | $\hat{\beta}$ | SE | p | $\hat{\beta}$ | SE | p | $\hat{\beta}$ | SE | p | $\hat{\beta}$ | SE | p | $\hat{\beta}$ | SE | p | $\hat{\beta}$ | SE | p | $\hat{\beta}$ | SE | p | $\hat{\beta}$ | SE | p | $\hat{\beta}$ | SE | p | $\hat{\beta}$ | SE | p |
| Intercept |  | 0.23 | 0.63 |  | 0.25 | 0.27 |  | -1.32 | 0.30 | *** | -1.83 | 0.60 | ** | -0.63 | 0.81 |  | -0.41 | 0.34 |  | -0.66 | 0.09 | *** | -0.69 | 0.12 | *** | -0.89 | 0.14 | *** | -1.56 | 0.25 | *** | -0.51 | 0.19 | ** | -0.43 | 0.44 |  |
| Dist.tree |  |  |  |  |  |  |  |  |  |  |  |  |  |  |  |  |  |  |  |  |  |  |  |  |  |  |  |  |  |  |  |  |  |  | 0.57 | 0.21 | ** |
| Tree.H |  |  |  |  |  |  |  |  |  |  | 0.72 | 0.31 | * |  |  |  |  |  |  |  |  |  | 0.18 | 0.09 | * | 0.32 | 0.09 | *** |  |  |  | 0.15 | 0.07 | * |  |  |  |
| Traffic |  |  |  |  |  |  |  |  |  |  |  |  |  | 2.79 | 1.81 |  |  |  |  | 0.79 | 0.09 | *** | 0.60 | 0.10 | *** | 0.53 | 0.12 | *** | 0.89 | 0.25 | *** | 0.65 | 0.08 | *** |  |  |  |
| Landscape type | DPT |  |  |  |  |  |  |  |  |  |  |  |  |  |  |  |  |  |  |  |  |  |  |  |  |  |  |  |  |  |  |  |  |  | 1.12 | 0.52 | * |
|  | FE |  |  |  |  |  |  |  |  |  |  |  |  |  |  |  |  |  |  |  |  |  |  |  |  |  |  |  |  |  |  |  |  |  | -0.18 | 0.80 |  |
|  | NV |  |  |  |  |  |  |  |  |  |  |  |  |  |  |  |  |  |  |  |  |  |  |  |  |  |  |  |  |  |  |  |  |  | 2.03 | 1.18 | . |
|  | PT |  |  |  |  |  |  |  |  |  |  |  |  |  |  |  |  |  |  |  |  |  |  |  |  |  |  |  |  |  |  |  |  |  | -0.27 | 0.72 |  |
|  | SPT |  |  |  |  |  |  |  |  |  |  |  |  |  |  |  |  |  |  |  |  |  |  |  |  |  |  |  |  |  |  |  |  |  | -0.16 | 0.55 |  |
| Julian Day |  | 0.74 | 0.53 |  | -0.40 | 0.59 |  | 0.49 | 0.19 | ** | -0.46 | 0.58 |  | -1.00 | 0.48 | * | -0.45 | 0.30 |  | -0.01 | 0.10 |  | 0.20 | 0.10 | . | 0.05 | 0.12 |  | -0.45 | 0.21 | * | 0.09 | 0.08 |  | -0.16 | 0.25 |  |
| Julian Day<sup>2</sup> |  | 0.10 | 0.74 |  | -1.00 | 0.58 | . | 0.37 | 0.17 | * | 0.49 | 0.45 |  | -0.10 | 0.38 |  | -0.16 | 0.24 |  | 0.07 | 0.09 |  | -0.11 | 0.10 |  | 0.08 | 0.11 |  | 0.23 | 0.18 |  | 0.02 | 0.07 |  | -0.10 | 0.19 |  |

p < 0.1 = . p < 0.05 = \* p < 0.01 = \*\* p < 0.001 = \*\*\*

**Figure 6:**
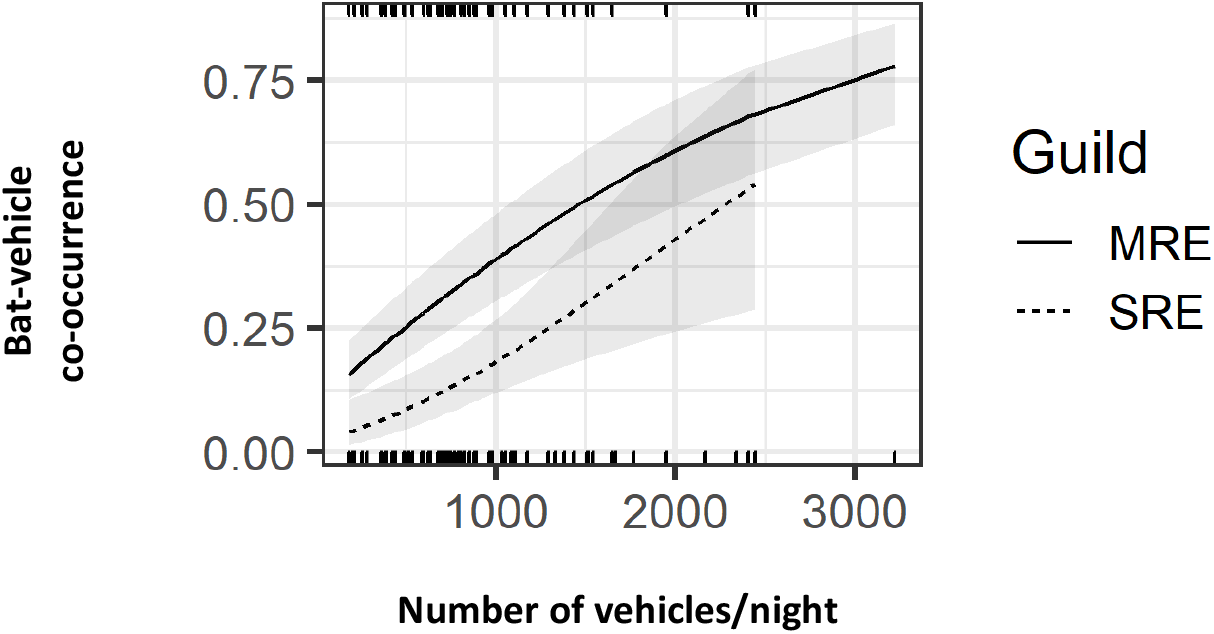
Predicted effect of traffic volume on proportion of bat-vehicle co-occurrence (trajectories positioned at risk and at more than 10 s from a vehicle pass). 95% confidence intervals are shown. Ticks in x axis represent sampled values (bottom = MRE; top = SRE). SRE: short-range echolocators. MRE: mid-range echolocators.

### Model 4 – Orientation of flight trajectories

The large majority of flight paths followed the road axis in all landscape types (**Figure A 3**). Landscape type, distance to tree foliage, and tree height were not selected to explain trajectory orientation (**Table 5**). Nonetheless, an increasing traffic volume was associated to a larger proportion of trajectories parallel to the road axis in *P. pipistrellus* (**Table 5**). Season had a very weak, or even no effect on the proportion of trajectories parallel to the road (**Figure A 2**).

**Table 5:**
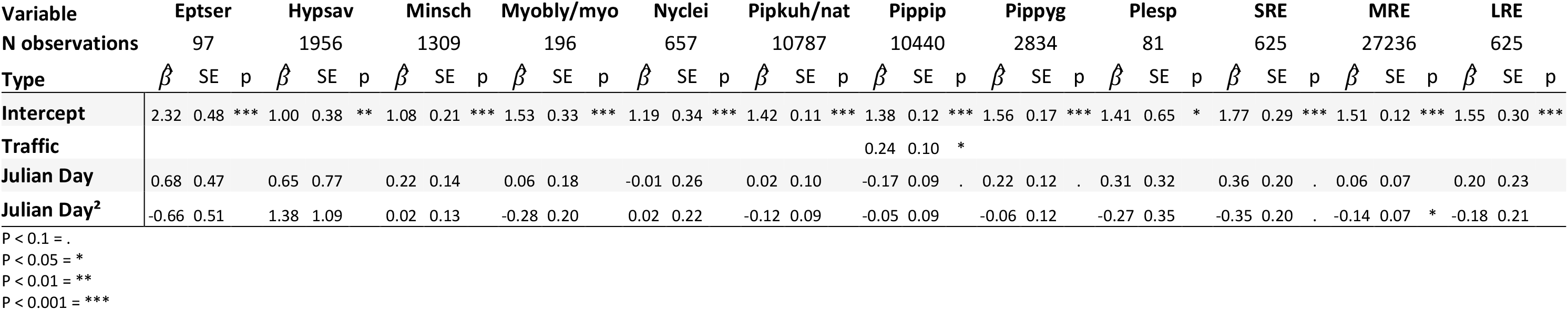
Summarised statistical results of the binomial distributed generalised linear mixed effect models (GLMM) for the flight path orientation for each species. 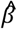 = estimate. SE = standard error. p = significance of p value. Species names are given with the three first letters of the species and genera. Dist.tree = distance to tree foliage. LRE: long-range echolocators. MRE: mid-range echolocators. SRE: short-range echolocators.

### Product: number of bat passes at collision risk per night

There was only a small selection of species for which the same variable had an effect on at least two aspects of the collision risk (i.e. on bat density, bat presence in the zone a risk, or bat-vehicle co-occurrence). These cases are all described in this section. An increase in traffic was associated with a tendency of an increasing number of bat passes at risk of collision for *P. kuhlii/nathusii* and the SRE guild (**Figure A 4**). The number of bat passes at collision risk was higher at perpendicular tree rows for *P. pipistrellus* but higher at forests and forest edges for *H. savii* (**Figure A 4**). The number of bat passes at risk of collision was higher in summer for *E. serotinus*, *H. savii*, *P. kuhlii/nathusii* and *P. pipistrellus*, while it was higher in autumn for *M. schreibersii*, *M. daubentonii*, *P. pygmaeus* and *Plecotus sp*, and higher in spring for *M. myotis/blythii* (**Figure A 2**).

We found a mean number of bat passes at risk of collision per kilometre and per night of 2.3 for SRE, 1024.9 for MRE and 11.7 for LRE (**Figure 7**). The index of susceptibility to road collisions, which is independent of species population densities, placed MRE as the most susceptible guild (**Figure 8**).

**Figure 7:**
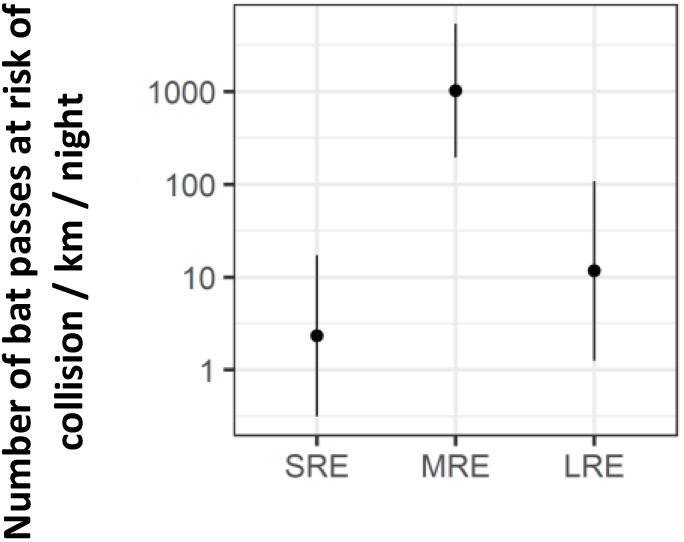
Predicted number of bat passes at, risk of collision per night and per kilometre for each bat guild (logarithmic scale). 95% confidence intervals are shown. SRE: short-range echolocators. MRE: mid-range, echolocators. LRE: Long-range echolocators.

**Figure 8:**
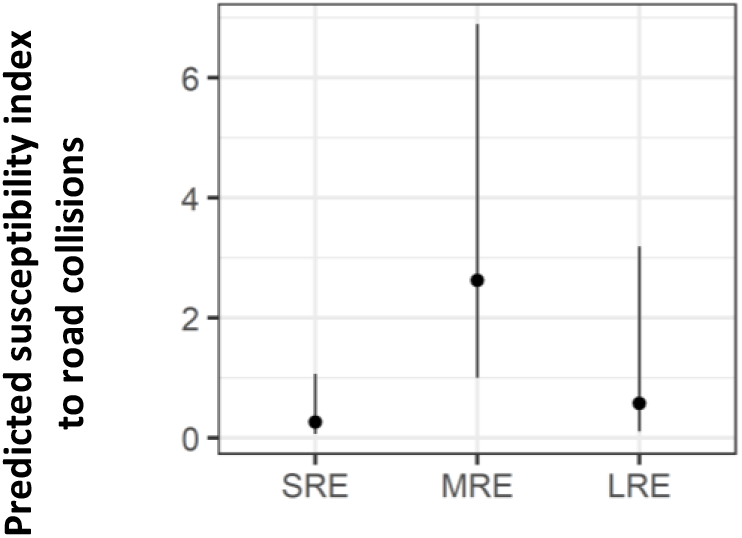
Predicted susceptibility index to road, collisions for each bat guild. 95% confidence intervals are shown. SRE: short-range echolocators. MRE: mid-range echolocators. LRE: Long-range echolocators.

## Discussion

This study aimed at disentangling the different mechanisms that influence bat-vehicle collision risks: (1) density of individuals recorded from the road edge, (2) position in the zone at risk (low flight height over the road), (3) bat-vehicle co-occurrence, and (4) flight path orientation. Disentangling those mechanisms was possible thanks to three-dimensional acoustic flight path tracking of bat behaviour at roads on a large scale (French Mediterranean region) on a large amount of data (122,294 bat passes). Our results demonstrate heterogeneity in the influence of explanatory variables on the four response variables, depending on species.

### Local landscape type

The effect of local landscape type on density was heterogeneous according to species. For example, perpendicular tree rows led to a higher density of *P. pygmaeus* compared to simple parallel tree rows, and forest edges led to a higher density of *H. savii* compared to simple parallel tree rows. It is rather unlikely that the structural composition (density and orientation of linear vegetation) is the only explanation for these differences, because other confounding effects may very well increase bat density also, such as the different tree species that were often associated with a type of landscape; for example, it is known that *P. pygmaeus* prefers riparian habitats (Rachwald et al., 2016).

Interestingly, landscape type did not produce similar effects on bat density and on bat flight behaviour. Indeed, in the case of *P. pipistrellus* for instance, bat density was the highest in perpendicular tree rows formed by small streams. However, local landscapes eliciting the highest proportion of flights at collision risk for this species were forests. This type of landscape was in fact a very high factor of presence at risk for most species and guilds, as we expected. Vegetation density may play an important role here. Indeed, parallel tree rows consisted in individual planted trees with gaps of 10 to 20 m while forests had dense shrub layers most of the time. Tree rows probably allow bats to benefit from the edge effect (i.e. easy access to flying insects abundant in or near tree foliage) (Brigham et al., 1997; Evans et al., 2003; Verboom and Spoelstra, 1999) without needing to fly directly above the road, contrarily to forest landscapes with hard edges, which act as conduits (Kalcounis-Rueppell et al., 2013).

Contrary to our expectations, landscape types were not selected to explain the orientation of flight trajectories. Our results show that even in the presence of a perpendicular tree row or in the absence of trees, bats fly most of the time parallel to the road axis. This supports the idea, not often enough emphasised in collision risk assessments, that bats may use roads as corridors, because road verges may offer foraging opportunities by attracting more insects than adjacent habitats (Medinas et al., 2019; Villemey et al., 2018), and because of the verge effect when trees are present (Brigham et al., 1997; Kalcounis-Rueppell et al., 2013; Verboom and Spoelstra, 1999). According to our results, it should be considered that on secondary roads, bats following the road axis may be as common as bats crossing roads, and that mitigation measures should deal with these two types of movements.

### Tree height and distance to tree foliage

As said in material and methods, distance to tree foliage was correlated with tree height in our study. In all species except *Plecotus sp*., density was negatively affected by an increasing distance to tree foliage, when selected. Conversely, taller trees led to a higher density of *M. myotis/blythii* and of the LRE guild. Our hypothesis according to which taller trees would be associated with a higher density was thus only verified for one species. The effect of distance to trees was shown in several studies out of road context (Heim et al., 2015; Kelm et al., 2014; Verboom and Spoelstra, 1999), and once at hedgerows crossing roads (Abbott, 2012), but never at road study sites with different landscape structures and for several distinctly identified species. In our study, the proportion of flights in the zone at collision risk was rarely influenced by tree height or distance to trees; nonetheless, increasing distance to trees was associated with higher proportions of flights at risk for *E. serotinus*, contrary to our expectations. In addition, in several species (*M. daubentonii, P. pipistrellus* and *P. pygmaeus*), taller trees (that generally had their foliage over the road) led to a higher bat-vehicle co-occurrence.

### Traffic

Traffic volume did not affect bat density or position in the zone at risk for most species, contrary to our expectations. Nonetheless, *P. kuhlii/nathusii* flew less in the zone at collision risk when traffic increased, possibly because they recognise the danger associated with vehicles, although a specific data set would be required to test this hypothesis. Moreover, *P. pipistrellus* flew parallel and over the external sides of the road more often with increasing traffic. These results complete the observations of Zurcher et al. (2010), who did not distinguish between species, but found that 60% of approaching individuals reversed their course in the presence of a vehicle.

Our results trivially showed that the higher the traffic volume, the higher the temporal bat-vehicle co-occurrence, probably because bats have no choice but to cross the road closely in time with vehicle passes when traffic is high. In addition, our study showed that SRE are less likely to fly in the zone at collision risk when a vehicle is present compared to MRE. Since the foraging abilities of SRE seem to be more impaired by light and noise than MRE (Azam et al., 2018; Siemers and Schaub, 2011; Stone et al., 2015), MRE might use roads as foraging grounds and take more risks than SRE. Therefore, even if SRE are known to fly lower than MRE and thus at heights more similar to those of vehicles (Berthinussen and Altringham, 2012; Roemer et al., 2019), their lower bat-vehicle co-occurrence should partially mitigate their susceptibility to collisions. This result emphasises the importance of accounting for the different aspects of species behaviour when evaluating their susceptibility to collisions (Chamberlain et al., 2006).

### Time of year

Our results show typical activity patterns throughout the year with peak density in summer or autumn, that seemingly drive the number of bat passes at collision risk per night (the product of quantitative models), that also shows a peak in summer or autumn. However, it is the first time to our knowledge that it is demonstrated that flight proportion in the zone at risk at roads increases in autumn (for several species and guilds). An increased flight proportion in the zone at risk in autumn could partly be attributed to the naïve behaviour of juveniles, which after birth and emancipation suddenly increase population sizes at the end of the summer (Dietz et al., 2009), and that are, necessarily present in our dataset even if we cannot assess their proportion. Juveniles are indeed more vulnerable to road collisions than adults (Fensome and Mathews, 2016). This result could also be explained by increasing foraging opportunities on roads during colder times, as was observed in swallows (Evans et al., 2003), and increased energetic demands before hibernation (Dietz et al., 2009).

### Species differences

Our study provides detailed information at the species level except for species with small sample sizes, for which readers are referred to the guild level. Models for guilds also inform on the extent of generalisation of results because variables selected at the guild level are assumed to exert a significant influence on several species.

Forests clearly stood out as a landscape type with a higher proportion of trajectories in the zone at risk for MRE, *P. pipistrellus* and *P. kuhlii/nathusii*. For *H. savii* and *E. serotinus*, double parallel tree rows elicited the smallest proportion of trajectories in the zone at risk. In most species, locations without trees generated a relatively low proportion of trajectories in the zone at risk.

We found a mean number of bat passes at risk of collision per kilometre and per night of 2.3 for SRE, 1024.9 for MRE and 11.7 for LRE. We stress that these figures are necessarily an overestimate since we could not measure more precisely bat avoidance of vehicles when they were in the zone at collision risk at less than 10 s from a vehicle pass. In addition, readers have to bear in mind that these figures are not a proxy for the bat guild susceptibility to road collisions. For this, it is necessary to consider the proportion of individuals in the zone at collision risk multiplied by the co-occurrence of bats and vehicles. This calculus placed MRE as the most susceptible bat guild to road collisions. This finding did not match our expectations since the lowest flyers were always thought to be the most susceptible to road collisions (Voigt and Kingston, 2016). Fensome and Mathews (2016) found that low-flying species are more susceptible to collisions, however, it is important to mention that they included both SRE and MRE in this category. Our results show that MRE are more susceptible than SRE to road collisions because MRE fly more often in the zone at collision risk and are also more often present in this zone simultaneously to a vehicle pass. This classification, added to species conservation status, can be used to prioritise conservation actions at roads.

### Advantages of conditional probabilities taking into account bat behaviour to assess road collision risks

All quantitative models succeeding the density model were interpreted as conditional probabilities that an individual is at risk of collision, and their predicted probabilities were multiplied to obtain the overall bat collision risk if a variable was selected in several of them. The product of all quantitative models showed that *H. savii* was more at risk of collision at forests and forests edges (and to a lesser extent at roads without trees), while *P. pipistrellus* was more at risk of collision at perpendicular tree rows. These products match the patterns of bat density in function of landscape type. The product of quantitative models also showed that the yearly patterns of collision risks matched the ones of bat density. Collision risks are more numerous in summer or autumn according to species, and explain the mortality patterns found in Fensome and Mathews (2016).

However, while increasing traffic density was associated with a decrease in SRE density, it was associated with an increase in the overall collision risk (the product of quantitative models). This demonstrates, as we expected, that the measure of the number of bat passes can be a good proxy of, bat collision risks in certain contexts, but that it is necessary to also measure bat behaviour to assess collision risks with certainty in all contexts.

In addition, contrary to the classic method of collecting bat carcasses, the results of acoustic flight path tracking are not biased by predation or observer efficiency, and acoustic flight path tracking may be applied to study roads after as well as before they are in service, if necessary. It is also well known that bat carcasses are quite difficult to find (Santos et al., 2011; Slater, 2002) while acoustic flight path tracking provides a large amount of precise information on bat movements. Yet, out of curiosity, during field work, we looked for bat carcasses at least once per study site, most often twice (on two different days), and more rarely up to four times (on four different days). Searches were done along the road on sections 50 m in length, on each side of the study point. Because it was not the purpose of our study, searches were randomly done during the day (from 9 am to 9 pm), which has an influence on the finding success since small carcasses are rapidly scavenged (Santos et al., 2011; Slater, 2002). Nevertheless, only 2 carcasses were found overall (unpublished data). One juvenile female of *Rhinolophus hipposideros* was found on the 12th of August 2016 at study site #11 (dense oak forest on both sides) and one adult *Pipistrellus pipistrellus* was found on the 7th of June 2016 on study site #55 (“no vegetation”: some vines and croplands). These results underline the fact that to attain the aims of our study and to collect enough data per species with direct counts of bat carcasses, it would have been necessary to invest a significantly greater amount of time than it was necessary using acoustic recordings.

### Recommendations for road siting and management

Our first group of recommendations applies to habitat selection during road planning to avoid situations with enhanced collision risks. As has been recommended in previous studies (Fensome and Mathews, 2016; Medinas et al., 2013), ‘quality habitats’ – depending on species ecology – should generally be avoided to ensure that roads will avoid habitats with high bat density. However, bat activity is highly dependent on distance to roost and may be under- or overrepresented at certain habitats according to the distance to roosts (Rainho and Palmeirim, 2011). Therefore, measuring species activity at different seasons on site will always provide more insights on the potential risks. Moreover, since it is assumed that species do not have comparable susceptibilities to road collisions (Fensome and Mathews, 2016), possessing information on species presence and density is highly relevant. The present study also allows us to emit recommendations for road siting based on the behavioural reactions to landscape features that we measured. Forested areas should be avoided because they elicit high proportions of flights at risk. Areas without trees should be prioritised because they almost always led to very low activity levels and low proportions of flights at risk. However, to explain position at risk, landscape types were only selected in models for species belonging to MRE and LRE and we cannot conclude on their effect on SRE.

Our second recommendation applies to the management of roadside vegetation during construction work and during the operational phase, to reduce collision risks. A gap of five meters between the road edge and tree foliage significantly decreased the activity levels of several species across the three different guilds. Our appreciation of study sites suggests that this effect could be due to higher primary productivity when vegetation is higher and closer to the road. If less primary biomass is available to insects, foraging opportunities for bats decrease, and so does their density (Threlfall et al., 2012). It is however controversial to recommend cutting trees at road sides, because this decision will engender habitat loss for numerous taxa, especially in large-scale impacted areas such as linear transport infrastructures. Opening habitat at road edges also creates suitable foraging grounds for birds of prey for instance (Morelli et al., 2014), and will increase their collision probability. It is possible to make these open verges less attractive by converting them to gravel surface (Kociolek et al., 2015), but this will eliminate plant habitats. In our results, hard edges also led to higher rates of MRE in the zone at collision risk. Another possibility for the management of vegetation is thus to only cut a certain number of trees and clear shrub layers periodically (a frequent practice in French Mediterranean forests to prevent fires) to reduce primary production and to allow bats to navigate between trees rather than above the asphalt. The local management will thus depend on the biodiversity stakes of the area. In areas of high stakes, reducing vehicle speed limit could be an efficient solution, but this was not tested on bats to our knowledge.

Finally, our results allow us to provide insight on a low-cost mitigation measure that has been popularly proposed to reduce collisions at secondary roads: hop-overs. They consist in planting tall trees at each side of a road to help bats increase their flight height and cross safely (Limpens et al., 2005). Screens can be added at each side of the road to prevent bats from crossing at low height. Christensen et al., (2016) already found that this measure could be ineffective to help many species crossing roads safely, as many individuals will just fly around screens to cross. Based on our results, we expect that planting tall trees next to roads will create new foraging grounds, increase bat density and encourage individuals to fly in the zone at collision risk if trees are planted very close to the road, as it is often recommended (Christensen et al., 2016; Voigt and Kingston, 2016). We therefore expect more collision risks with hop-overs than without, and their use without other measures such as speed reduction should be prohibited until their efficacy is proven.

### Limits of the study and perspectives

Our recommendations can only apply to landscapes and bat communities similar to the ones that we sampled. Therefore, complementary studies should be conducted in other biogeographical areas (e.g. Continental or Atlantic areas) to make sure that bats react consistently to the same road landscape features. However, we expect this endeavour to be quite difficult because of the local particularities in landscape management. Since we expect bats to be more active at prolific foraging grounds, it would be interesting to see if the measure of primary production - for example using satellite imaging - can be a more universal descriptor of bat activity than the description of the local landscape.

*Rhinolophus* species are assumed to be very susceptible to road collisions because they fly very close to ground level (Fensome and Mathews, 2016; Jones and Rayner, 1989; Roemer et al., 2017). However *Rhinolophus* species, because of their very high sonar frequencies (Kingston et al., 2000), are very difficult to detect and to record, and this is why we could not study their flight behaviour with our method. Acoustic flight path tracking with only two microphones would allow a study of *Rhinolophus* collision risks, although with simpler metrics (Claireau et al., 2018).

Several questions remain unanswered, such as the role of tree species, topography at a medium scale (i.e. slope of the terrain), and topography at a small scale (i.e. road embankments) in bat collision risks at roads. The nearby presence of a bat roost is also expected to be an important factor of, collisions. At last, it was reported that juveniles and males are more prone to road collisions (Fensome and Mathews, 2016). It would be interesting to study the behaviour of bats of different age and sex to explain this finding.

## Supplementary material

Script and codes are available online: https://github.com/Charlotte-Roemer/bat-road-collision-risks

Data are available online: https://doi.org/10.1101/2020.07.15.204115

## Acknowledgments

This study was a collaboration between Biotope and the Muséum national d’Histoire naturelle (Paris, France) in the form of a PhD thesis funded by Biotope and the Association Nationale de la Recherche et de la Technologie. We would like to thank Bruno Sanchez, Dominique Guicheteau (Réserve naturelle nationale de la Plaine des Maures), Ugo Schumpp and Julien Penvern for their precious help during field work. We thank Fiona Mathews for her comments before submission, as well as Gloriana Chaverri, Brock Fenton, Mark Brigham and two anonymous other reviewers for their comments. All of them greatly improved the quality of the manuscript. We are grateful to Kate Derrick for proofreading the manuscript. Version 3 of this preprint has been peer-reviewed and recommended by Peer Community In Ecology (https://doi.org/10.24072/pci.ecology.100067).

## Conflict of interest disclosure

Biotope is an environmental consultancy involved in road impact assessment studies. Two of the authors, Charlotte Roemer and Thierry Disca, were employees at Biotope during the time of the study. They thus declare a financial conflict of interest. Aurélie Coulon and Yves Bas declare that they have no financial conflict of interest with the content of this article. All of the authors take complete responsibility for the integrity of the data and the accuracy of their analysis. In addition, Aurélie Coulon is one of the PCI Ecology recommenders.

## APPENDIX

**Figure A 1:**
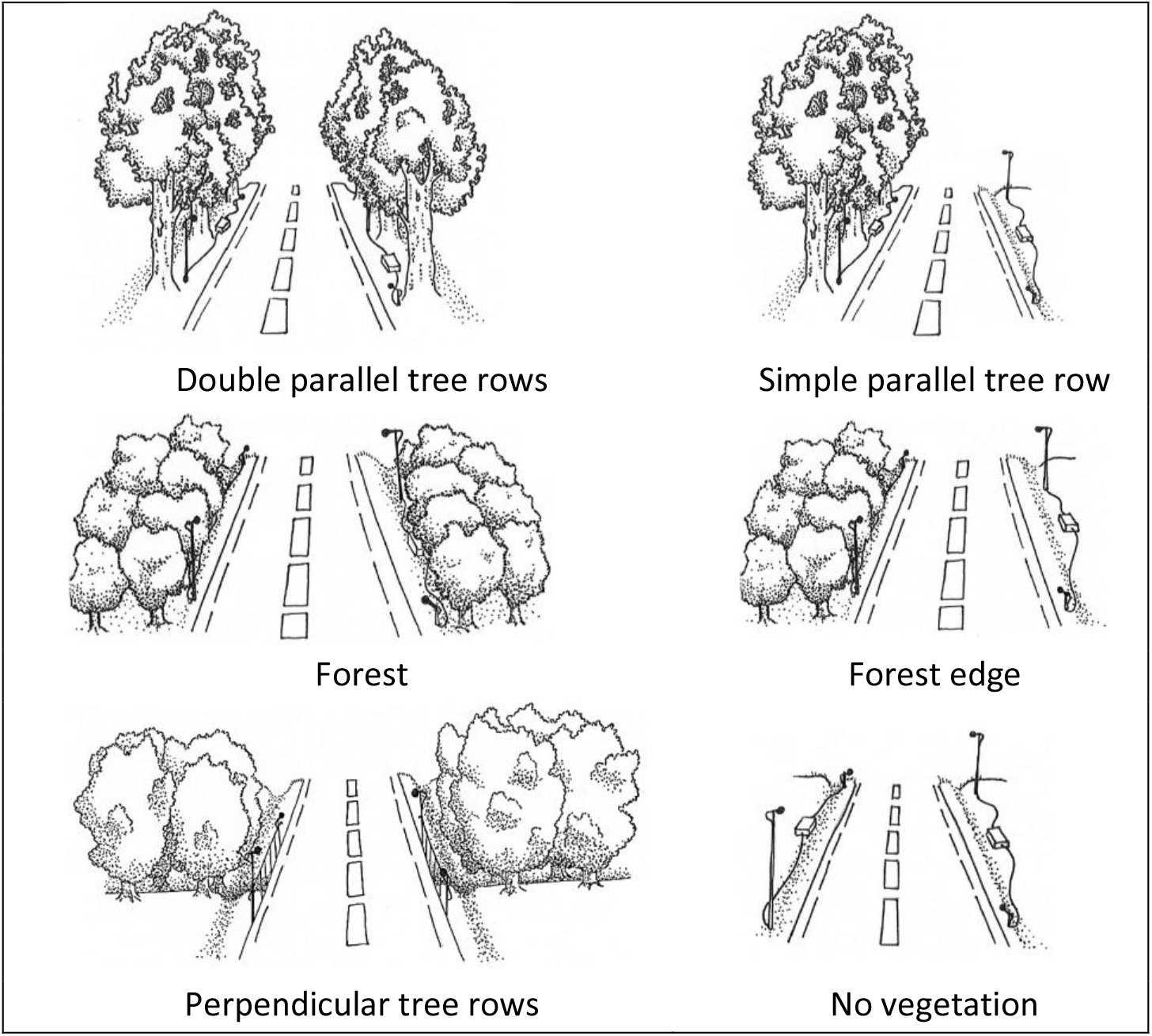
Landscape types. Microphones are shown on poles but when possible, they were attached to vegetation instead. The box between two microphones is the recorder.

**Table A 1:** Summary table for the total amount of data in each category and for each species. SRE: Short-range echolocators. MRE: Mid-range echolocators. LRE: Long-rang echolocators. N bat passes: rounded median number of bat passes among all four microphones. N flight trajectories: acoustic recordings with more than 3 consecutive calls recorded by all 4 microphones. N successful flight trajectories: flight trajectories with at least one call in the precision zone (<10 m from microphones centroid).

| Species | Guild | N bat passes | N flight trajectories | N successful flight trajectories |
| --- | --- | --- | --- | --- |
| *B. barbastellus* | SRE | 1157 | 311 | 97 |
| *M. alcathoe* | SRE | 2 | 1 | 1 |
| *M. bechsteinii* | SRE | 32 | 11 | 2 |
| *M. capaccinii* | SRE | 98 | 40 | 24 |
| *M. daubentonii* | SRE | 368 | 287 | 150 |
| *M. emarginatus* | SRE | 287 | 29 | 17 |
| *M. blythii/myotis* | SRE | 1477 | 583 | 201 |
| *M. nattereri* | SRE | 345 | 75 | 37 |
| *Myotis sp.* | SRE | 405 | 101 | 40 |
| *Plecotus sp.* | SRE | 908 | 281 | 89 |
| *R. euryale* | SRE | 6 | 0 | 0 |
| *R. ferrumequinum* | SRE | 68 | 2 | 2 |
| *R. hipposideros* | SRE | 129 | 3 | 1 |
| *H. savii* | MRE | 4634 | 3912 | 2053 |
| *M. schreibersii* | MRE | 9226 | 2268 | 1417 |
| *P. kuhlii/nathusii* | MRE | 45152 | 28601 | 11685 |
| *P. pipistrellus* | MRE | 32383 | 19830 | 11261 |
| *P. pygmaeus* | MRE | 14410 | 6136 | 3033 |
| *Eptesicus/Nyctalus/Vespertilio* | LRE | 503 | 141 | 31 |
| *E. serotinus* | LRE | 1161 | 475 | 107 |
| *N. lasiopterus* | LRE | 13 | 2 | 0 |
| *N. leisleri* | LRE | 9306 | 3506 | 698 |
| *N. noctula* | LRE | 32 | 16 | 5 |
| *T. teniotis* | LRE | 192 | 53 | 3 |
| TOTAL |  | 122294 | 66664 | 30954 |

**Figure A 2:**
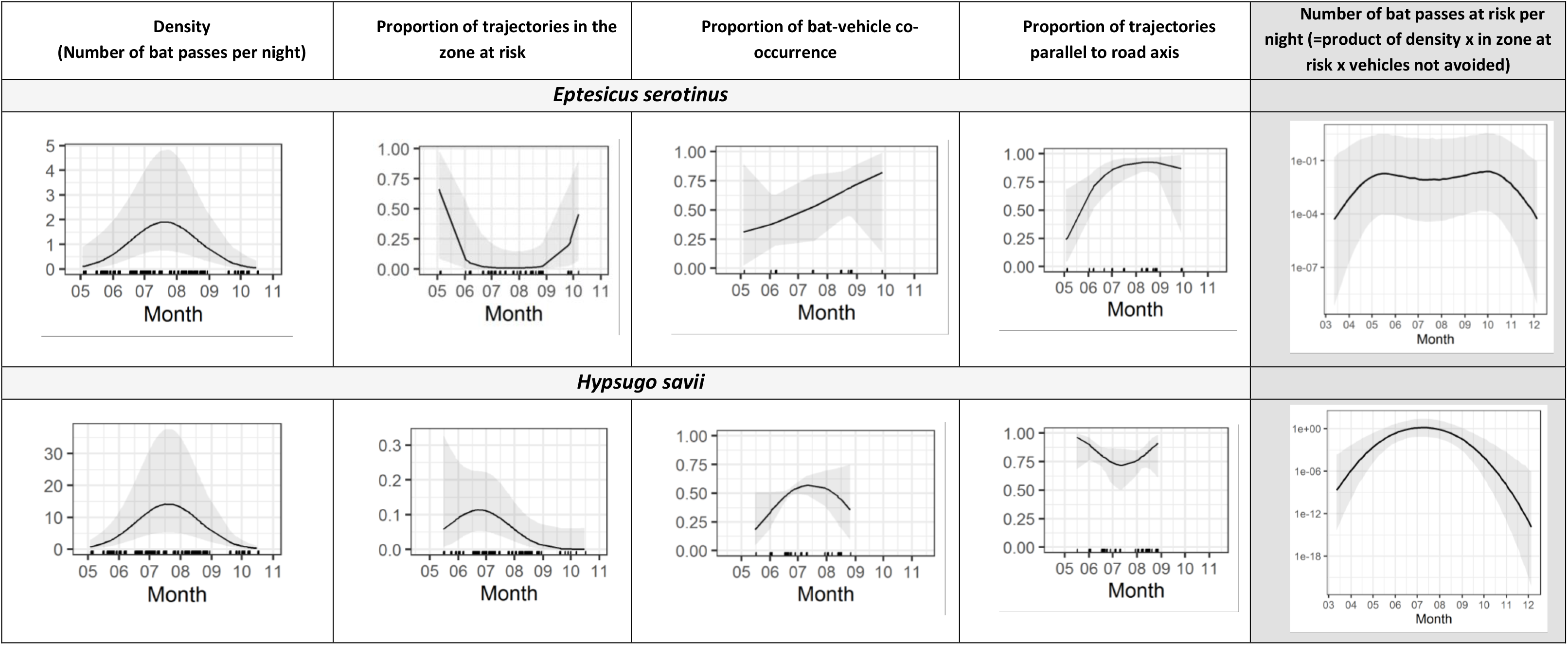
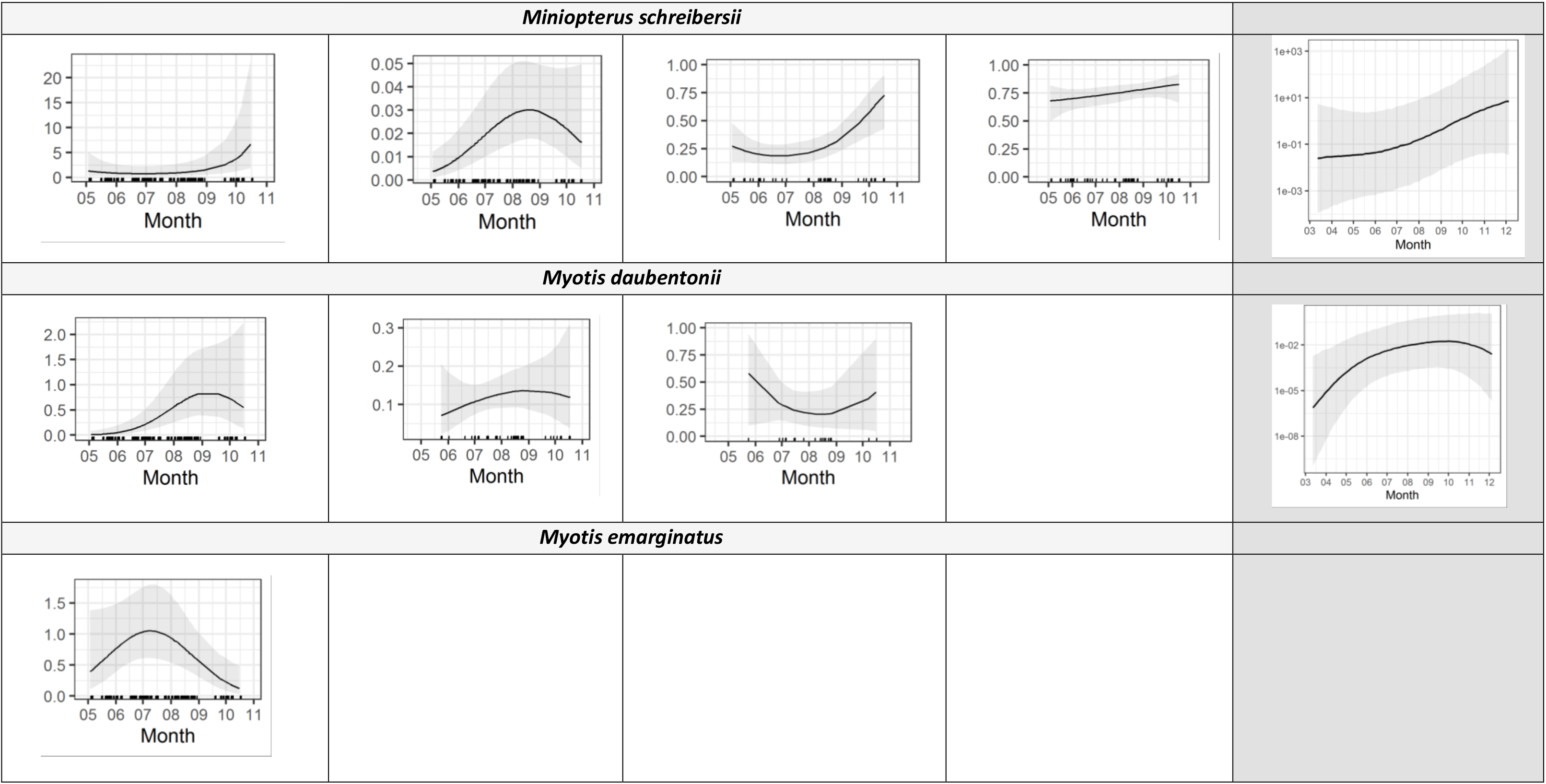
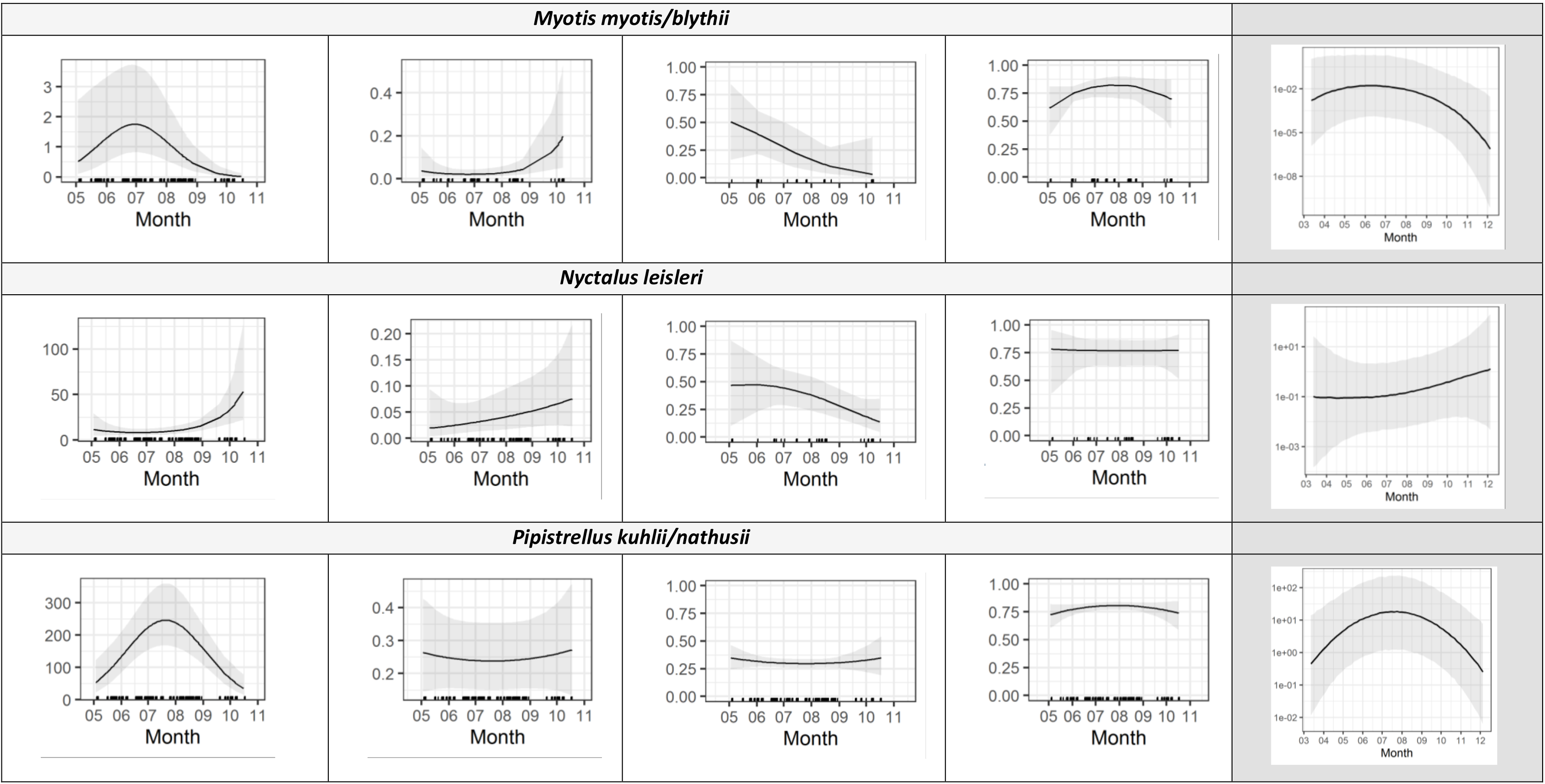
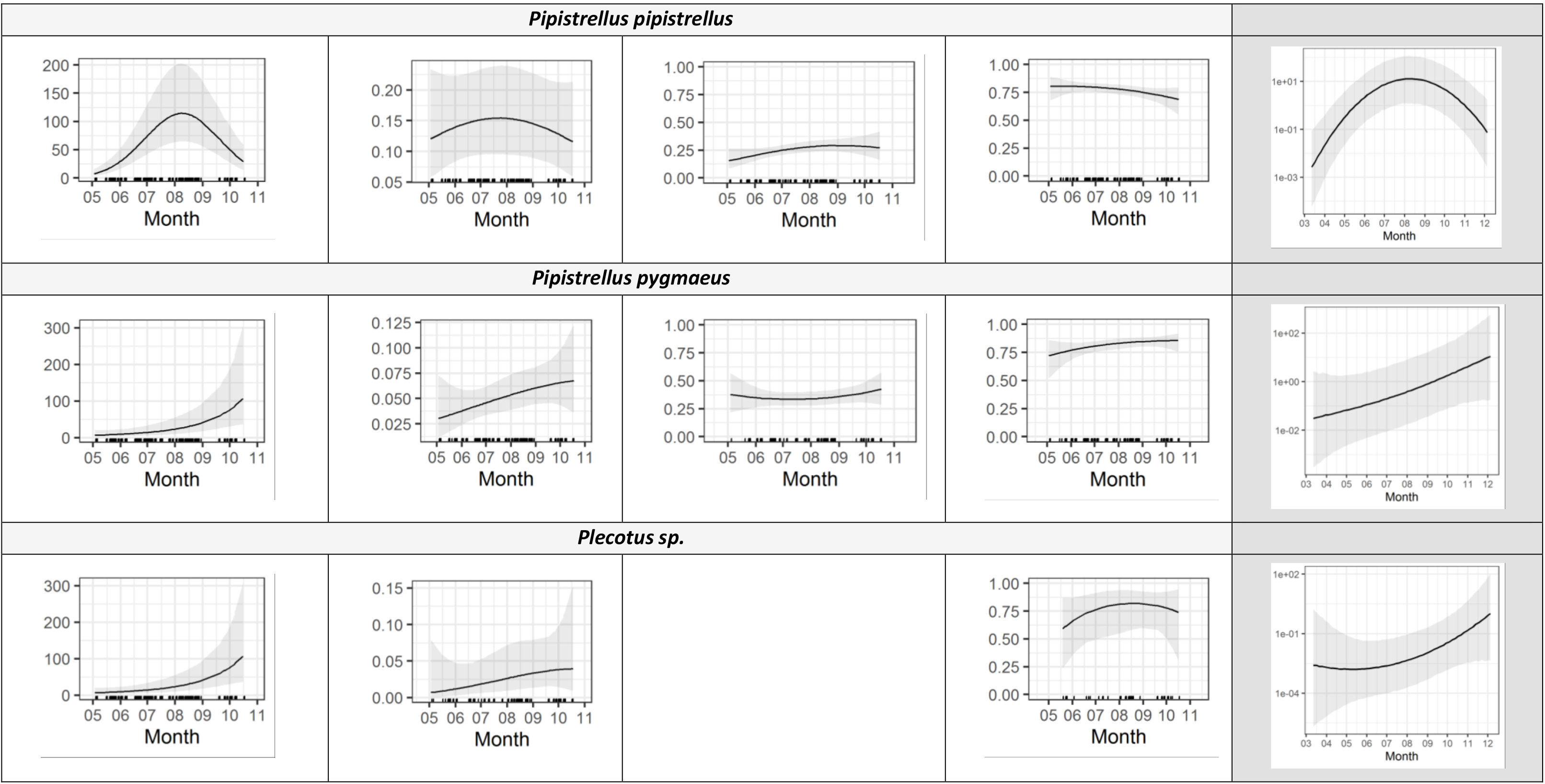
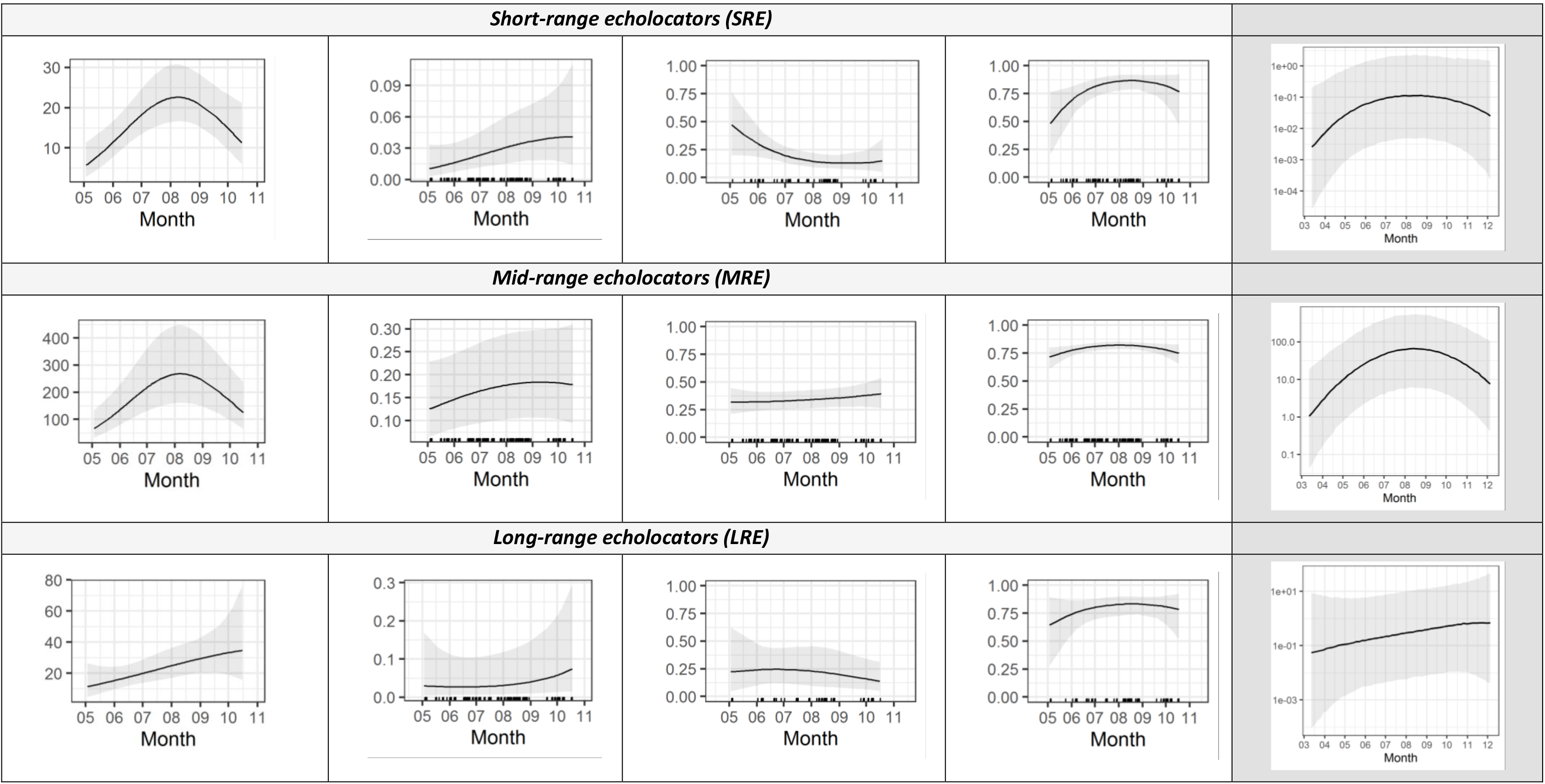
Predicted values for each model and each species or guild in function of time of the year (in month). 95% confidence intervals are shown. The last column displays the product of the first three models (density * in zone at risk * bat-vehicle co-occurrence) to obtain the number of bat passes at risk per night (logarithmic scale). Ticks on the x axis stand for the sampled values.

**Figure A 3:**
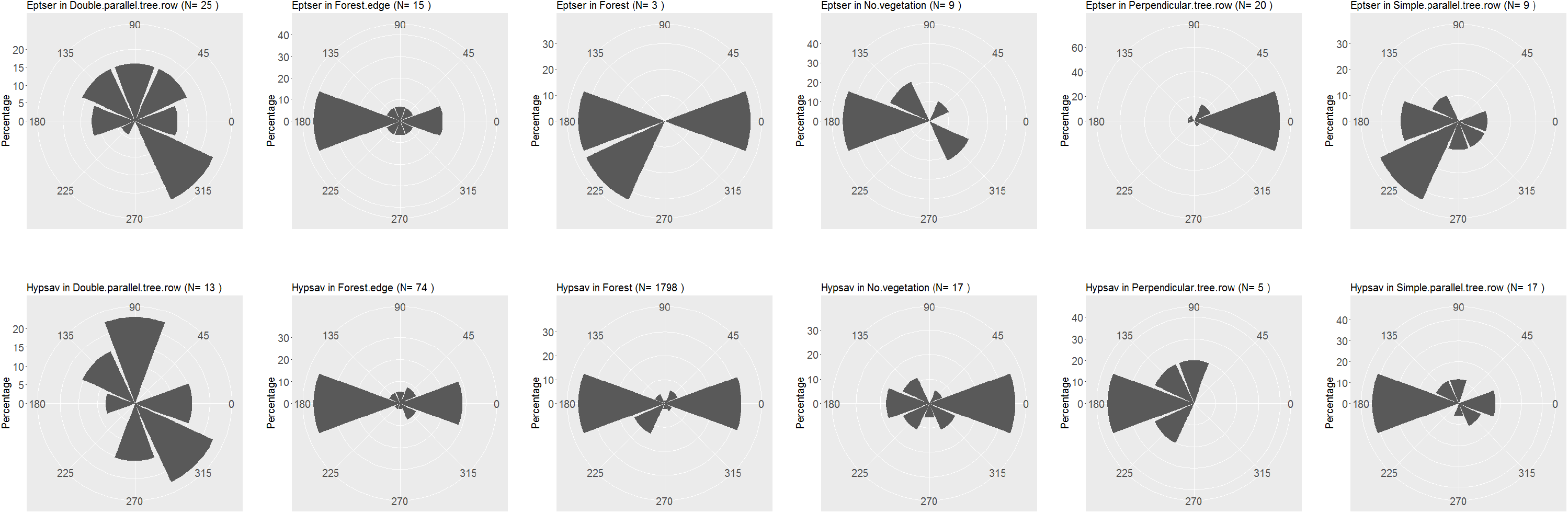
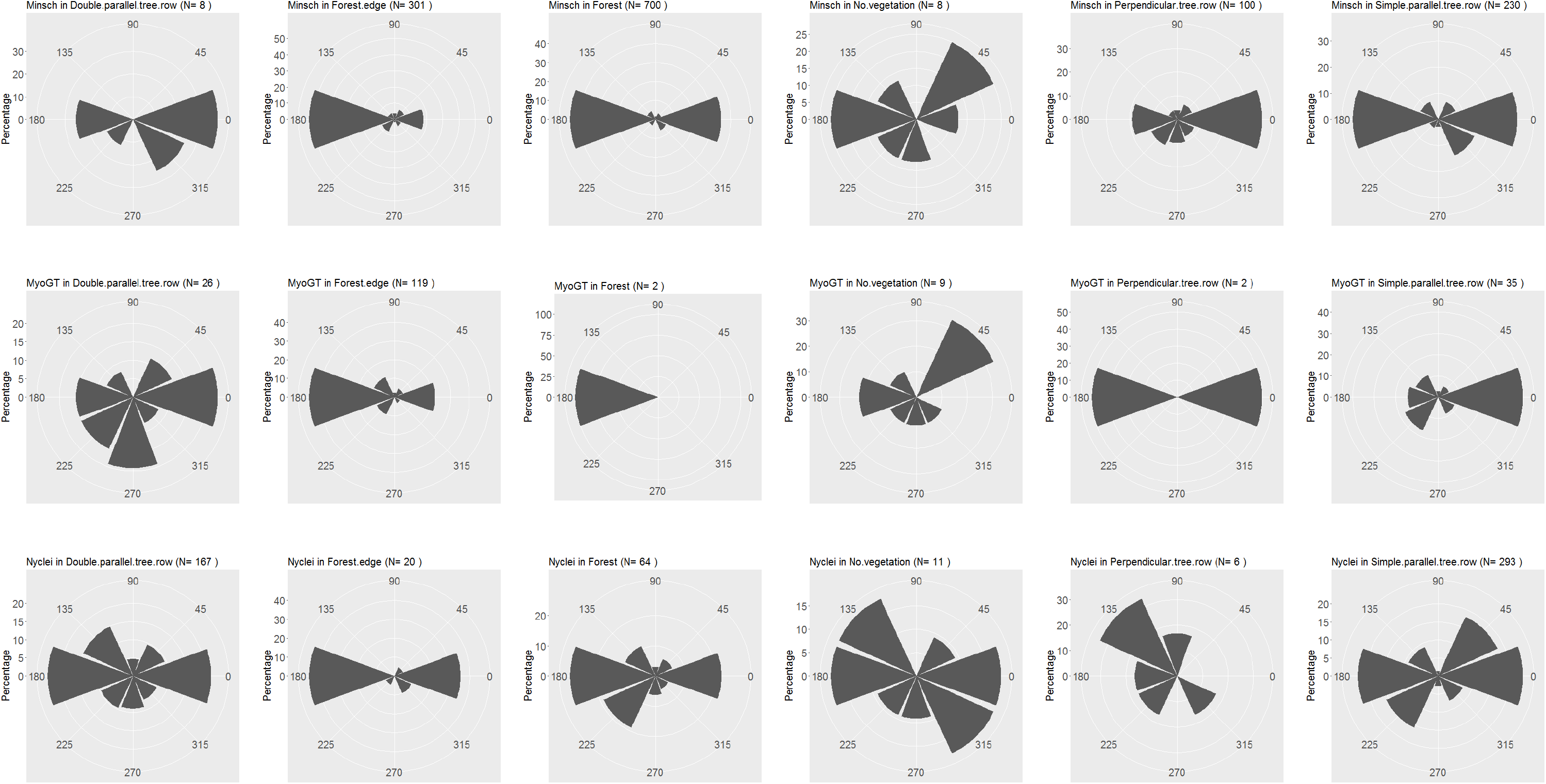
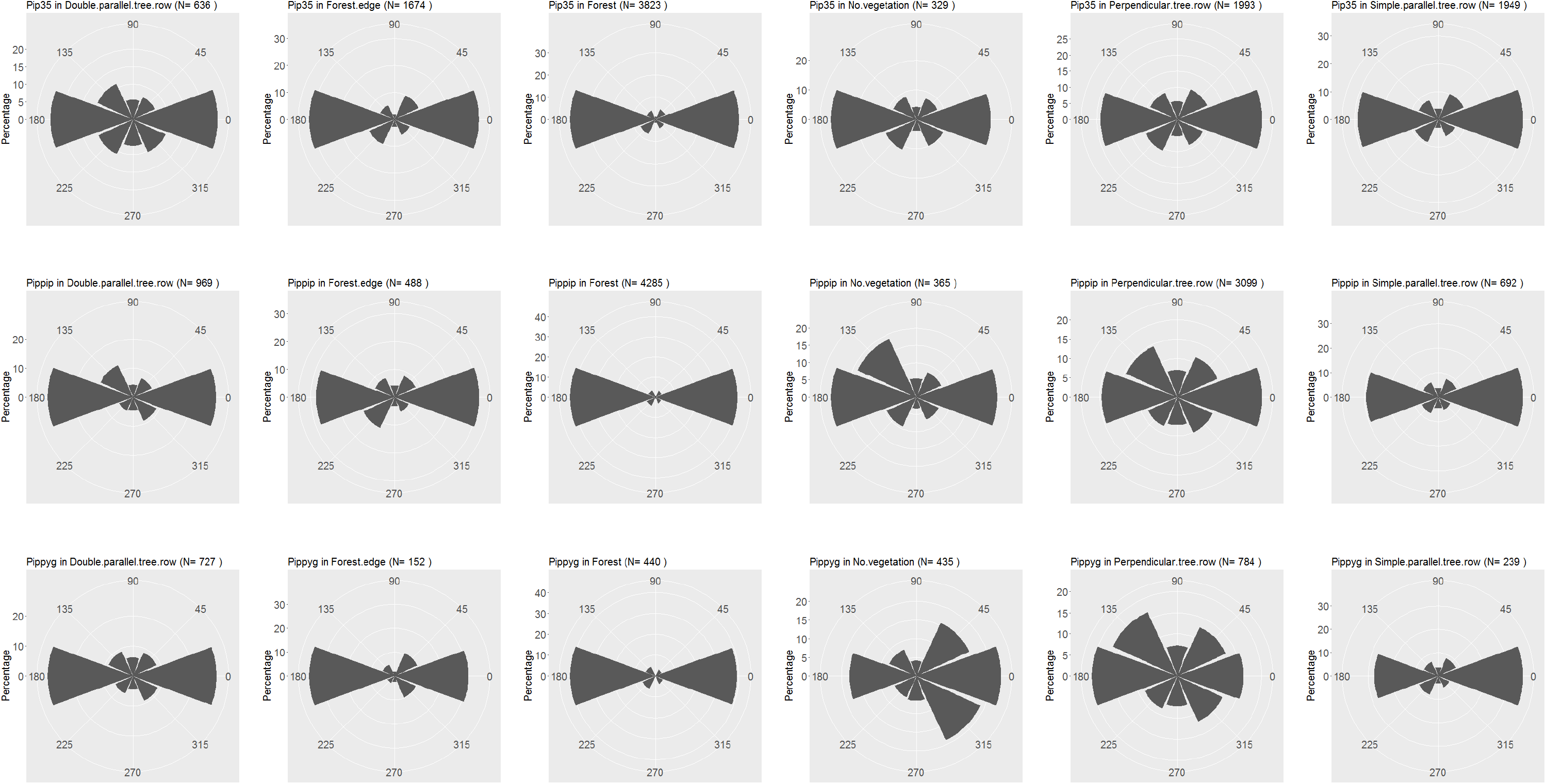
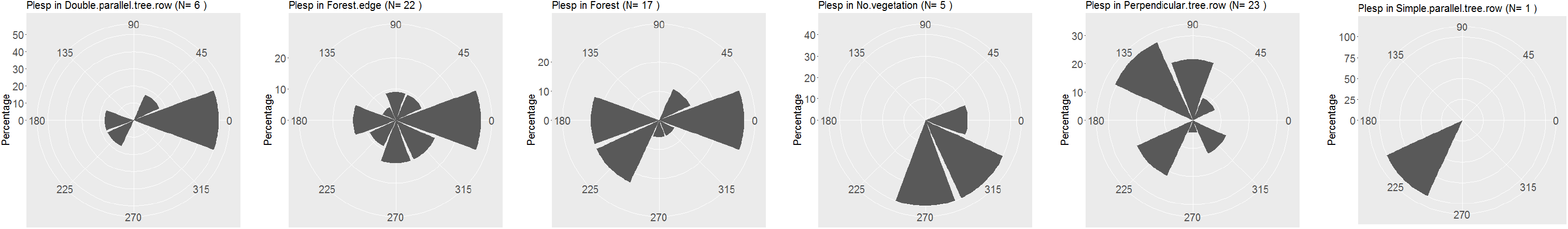
Polar representation of the orientation of bat flight paths in function of the landscape type (raw data). The road axis is represented by the axis 0° to 180°. N represents the number of flight paths that could be successfully located. Species names are given with the three first letters of the species and genera. The left axis provides the size of the scale in percentage of flight paths (i.e. for *E. serotinus*. in forests, 40 % of the flight paths are oriented toward 180°).

**Figure A 4:**
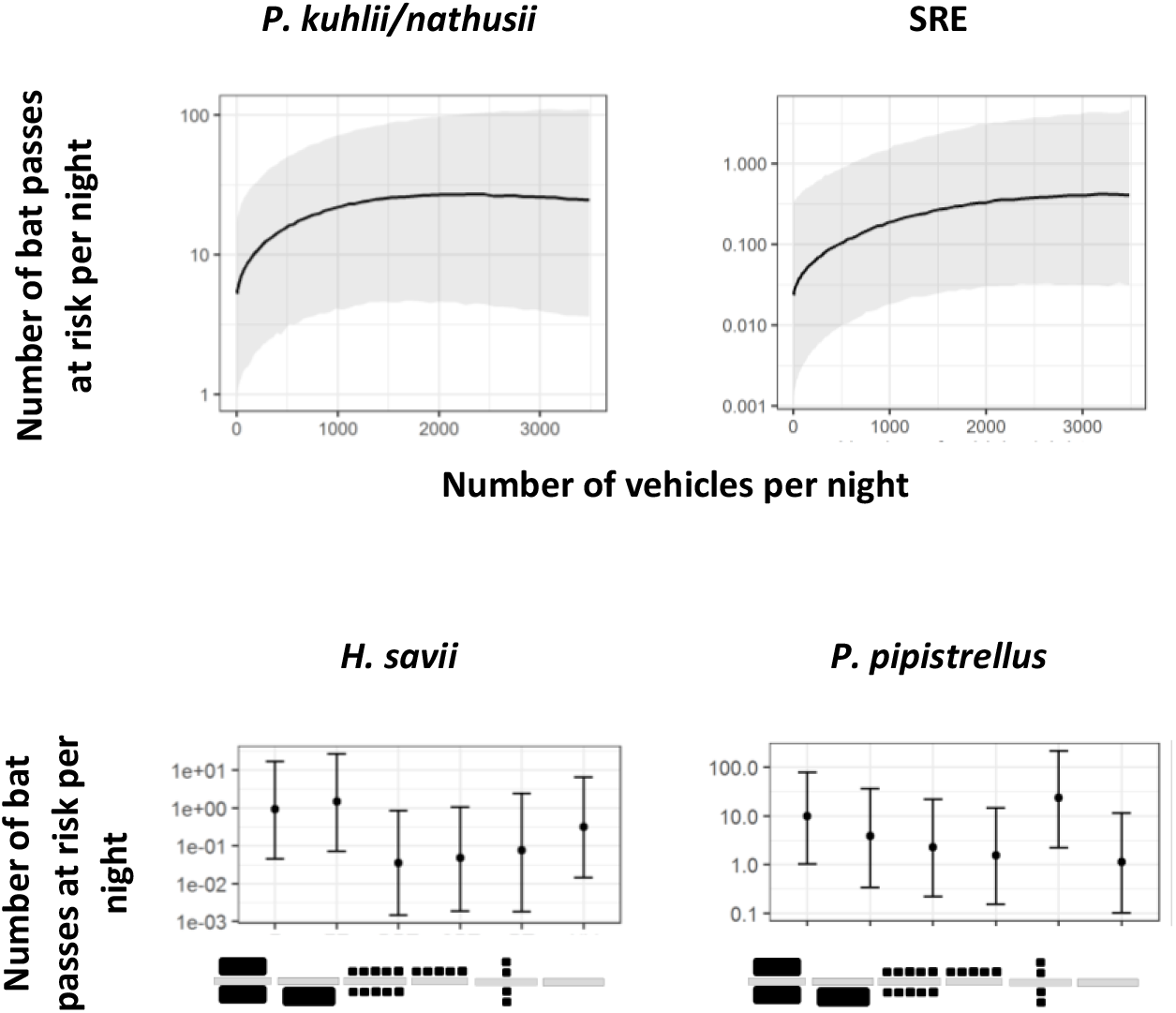
Product of the first three models (density * in zone at risk * bat-vehicle co-occurrence) to obtain the number of bat passes at risk per night (logarithmic scale) for variables selected in at least two models for the same species. 95% confidence intervals are shown. Figures represent landscape type viewed from the top (road in light grey and trees in black). SRE: short-range echolocators. LRE: long-range echolocators. MRE: mid-range echolocators. F = forest. DPT = double parallel tree rows. NV = no vegetation. FE = forest Edge. PT = perpendicular tree rows. SPT = simple parallel tree rows.

## Notes

https://github.com/Charlotte-Roemer/bat-road-collision-risks

### Summary of Updates

Version 3 of this preprint has been peer-reviewed and recommended by Peer Community In Ecology (https://doi.org/10.24072/pci.ecology.100067)

